# Development of digital Hi-C assay

**DOI:** 10.1101/2022.12.31.522397

**Authors:** Akihiro Mori, Gabriele Schweikert

## Abstract

**Abstracts**

Enhancers are genomic elements and contain all necessary cis-regulatory contexts. Such enhancers are convened to the appropriate promoter of target genes for gene regulations even though the enhancers and the promoters are apart a few mega-base pairs away from each other. In addition to physical distance, nucleotide mutations in enhancers influence a partial group of the target genes. Those make it more complicated to reveal the paired relationship between enhancer and promoter of target genes. Recently, advanced computational approaches are employed to predict such interactions. One approach requires a large number of different high-throughput datasets to predict such interactions; however, in practical aspects, all datasets for tissues and conditions of interest are not available. Whereas the alternative approach requires only genome sequences for particular predictions, their predictions are insufficient for practical applications. We address those issues by developing the digital Hi-C assay with a transformer-algorithm basis. This assay allows us to create models from simple/small/limited sequence-based datasets only. We apply the trained models to be able to identify long-distance interactions of genomic loci and three-dimensional (3D) genomic architectures in any other tissue/cell datasets; additionally, we demonstrated the predictions of genomic contexts by analysing the prediction patterns around the target locus in the three following genomic-context problems: enhancer-promoter interactions (i.e., promoter-capture Hi-C), the CTCF-enriched regions, and TAD-boundary regions. Because our approach adopted a sequence-based approach, we can predict the long-distance interactions of genomic loci by using the genomic sequences of the user’s interest (e.g., input sequences from high-throughput assay datasets such as ATAC-seq and ChIP-seq assays). Consequently, we provide an opportunity to predict interactions of genomic loci from a minimum dataset.

## Introduction

Enhancers are genomic elements and are often encountered outside the endogenous genomic context of their nearest genes^1^. The sequences of enhancers contain all necessary cis-regulatory contexts such as transcription factor binding sites (TFBSs). By using those contexts, the enhancers are convened to the appropriate promoter of target genes closely even though the enhancer and the promoter are apart a few mega-base pairs away from each other^2, 3^. Those enhancers’ functions and features can be altered by single nucleotide polymorphism (SNP) and indel mutations in the enhancer sequences significantly, resulting in leading to dysregulated target genes and developmental defects and causing a wide range of diseases^4–9^. However, such mutations influence a partial group of the target genes, which makes it more complicated to reveal the paired relationship between enhancer and promoter of target genes. Thus, it is important, but still largely unknown the general rules of controlling chromosome contacts and long-distance interactions.

With developing the latest sequencing technologies, we can identify the long-distance interactions of genomic loci such as enhancer-promoter interactions (EPIs) by conducting multiple high-throughput assays and integrating their results. For instance, DNase-seq and ATAC-seq assays can identify the open/active regulatory regions in chromosomes^10–12^. Chromatin immunoprecipitation (ChIP) with sequencing technologies^13^ can identify the genomic regions having specific histone markers. Among such histone markers, for example, H3K27ac and H3K4me1 correlate with the activations of transcription and promoters, respectively. Integrating these results allows us to narrow down the genomic locations of active enhancers and promoters in a specific condition (e.g., tissue-/cell-type, chemical treatments). Furthermore, capturing chromosome conformation (3C) technologies and their variants including Hi-C^14^ and ChIA-PET^15, 16^ allow us to identify the long-distance interactions of genomic loci comprehensively^17^. Integrating those experimental results allows us to identify the long-distance interactions of genomic loci including enhancer-promoter interactions. More recently, a variant and a highly specialised Hi-C, promoter-capture Hi-C (PCHi-C) has been introduced and applied to capture the interactions of more specific genomic loci (e.g., between promoter and enhancers regions)^18, 19^.

Despite these successes, those experimental approaches still have shortcomings. Firstly, it often overlooks a massive number of interactions. A single Hi-C and its variant approaches can capture millions of interactions but detect only 10-15% of genuine interactions (i.e., vital interactions with high false negatives) which hampers the identification of true positive interactions^20^. Secondly, those contemporary strategies require conducting multiple different types of high-throughput assays. Each assay result contains human and technical errors and biases; thus, the combined results potentially lose true positives, especially the ones in the long-range distance which rarely happens. Thirdly, due to technical difficulties including N-bp-cutter restriction enzymes, the resolutions of interactions are still as low as 1,000bp or longer if genome-scale research is conducted. A variant technology, Micro-C, overcomes this issue and allows us to access shorter-range interactions (between 200 bp and 4kb) at high resolution^21^. However, this approach tends to capture at most ∼10% of interactions even in their sample preparations (i.e., crosslinking), which again falls in the first shortcoming mentioned above. These technical drawbacks prevent identifying the interactions of genomic loci at a great distance, thus, hardly addressing to reveal the regulatory mechanisms of gene regulations and dysregulated genes causing medical diseases or developmental issues.

To complement these difficulties and to identify the long-distance interactions of genomic loci, especially enhancer-promoter interactions (EPIs), many advanced statistical and computational approaches including machine-/deep-learning algorithms have been applied^22–28^. As an advantage of those approaches, they allow us to train accurate models from raw data without prior knowledge; additionally, we can use these trained models to apply to a similar type of data to interpret the learned features and rules. Those approaches can be categorized into (1) genomic feature approach^26, 27, 29–36^ and (2) sequence-based approach^25, 37–40^. The genomic feature approach accepts similar concepts for determining EPIs experimentally and requires a large number of different high-throughput datasets to create prediction models, resulting in better prediction performance. In experimental aspects, however, all datasets for tissues and conditions of interest are not always available; hence, the datasets requirements can be a bottleneck of this approach^26, 27, 35, 41^. The sequence-based approach, whereas, aimed to achieve that the information in genome sequences within enhancers and promoters alone may be sufficient to distinguish EPIs. To do this, they require only a relatively simple and small pre-defined dataset (e.g., defined EPIs from Hi-C data) to create models^25, 38, 42^. However, their standard datasets contained largely overlapped regions; it has been reported that this approach performed poorly on cleaner datasets which removed overlapped regions or duplicated lines^43^. Nonetheless, both approaches achieved some success; thus, it is clear that signals from functional genomic data are informative in computationally distinguishing EPIs from non-interacting enhancer-promoter pairs.

With aiming an application in experimental aspects, we address the following question: can we develop a new computational approach which allows us to create models from simple/small/limited sequence-based datasets only and apply the trained models to identify EPIs and three-dimensional (3D) genomic architectures in any other tissue/cell datasets? To conduct this, we employ the sequence-based approach and address the shortcomings of this approach by adapting the several following conditions. Firstly, unlike the previous studies, we use shorter input sequences of a few hundred base pairs (bp), each of which is considered a representative region of a larger genomic region (e.g., a few thousand bp). Secondly, we do not predefine the enhancer or promoter regions. Instead, we use all raw interactions captured by a single Hi-C assay. Thirdly, we apply the custom one-chromosome-leave-out rule in which we train a model with all chromosomes leaving one chromosome out (i.e., inter-chromosome interactions) and then use the trained models to predict the interactions on the removed chromosome (i.e., intra-chromosome interactions) (material and methods). Finally, we employ the transformer algorithm^44^ instead of a recurrent neural network such as Long short-term memory (LSTM)^45^. Unlike the LSTM method, the transformer has an attention process: output at the decoder will try to look into multiple sections of the encoder. Since the transformer is a popular natural language processing for automated translating languages, we consider the interactions of genomic loci as a two-language relationship; a genomic locus (e.g., enhancer) as the input language and another genomic locus (e.g., promoter) as the output language.

Here we introduce the digital Hi-C assay with a transformer-algorithm basis. This assay requires only a single Hi-C data to predict comprehensive and long-distance interactions (i.e., equivalent to the interactions 10 times experimental assays). As a model case, we demonstrated this by training the models based on all raw interactions of the Hi-C data^20^ with the custom one-chromosome-leave-out rule and applying the trained model to predict interactions on the removed chromosome. We found our approach could predict 80-90% of interactions captured by cell-line specific Hi-C. By applying the trained models, further, we showed that our digital assay could predict the long-distance interactions in the same tissue/cell and different tissues. Furthermore, we showed that our approach could detect genomic context by characterising the patterns of overlooked (e.g., not predicting) interactions around the neighbour regions of the target locus. We demonstrated this on the datasets of three following genomic-context problems; enhancer-promoter interactions (i.e., PCHi-C)^46^, the CTCF-enriched regions^47^, and TAD-boundary regions^20, 48^. Because our approach adopted a sequence-based approach, we could predict the long-distance interactions of genomic loci by accepting input sequences from other high-throughput assay datasets such as ATAC-seq and ChIP-seq assays or even the sequence of the user’s interest. Consequently, we showed that our approach could provide an opportunity to predict interactions of genomic loci from a small dataset.

## Materials and Methods

### Data collection and vertical search approach to predict by transformer algorithm

All data are downloadable from either NCBI or European Nucleotide Archive (ENA) (figure 1 process 0). In the case of *.hic format, sequences are generated with Knight-Ruiz (KR) normalization^49^ with 5,000bp resolution. In the case of paired sequence format (i.e., *seq1 and 2), the KR normalization is applied. The sequence information is converted into genomic loci information and bins the interaction positions by 5,000bp (figure 1a process 1). The genomic versions were used originally cited. A 300 bp-length sequence is selected from the middle of each 5,000bp region (i.e., locus) and considered as a representative sequence for the region (figure 1a process 1). If highly similar sequences (i.e., sequence homology >90%) exist in the same dataset, new 300bp-length sequence candidates are re-selected randomly and checked for sequence similarity with other sequences in the same dataset. Repeat this process per re-selected sequence until all sequences in the dataset are not satisfied with the criteria of sequence similarity. To reduce the computational process, those sequences were converted into ‘N’-mer elements which were separated by a space (e.g., “acgtgctagc…” is converted to “acgtgcta cgtgctag gtgctagc …”). Note the N should be the same number as those of the ‘N’-mer token. In our study, N = 8 was set as a default condition (figure 1b-d, supplemental figure 1).

**Figure 1:**
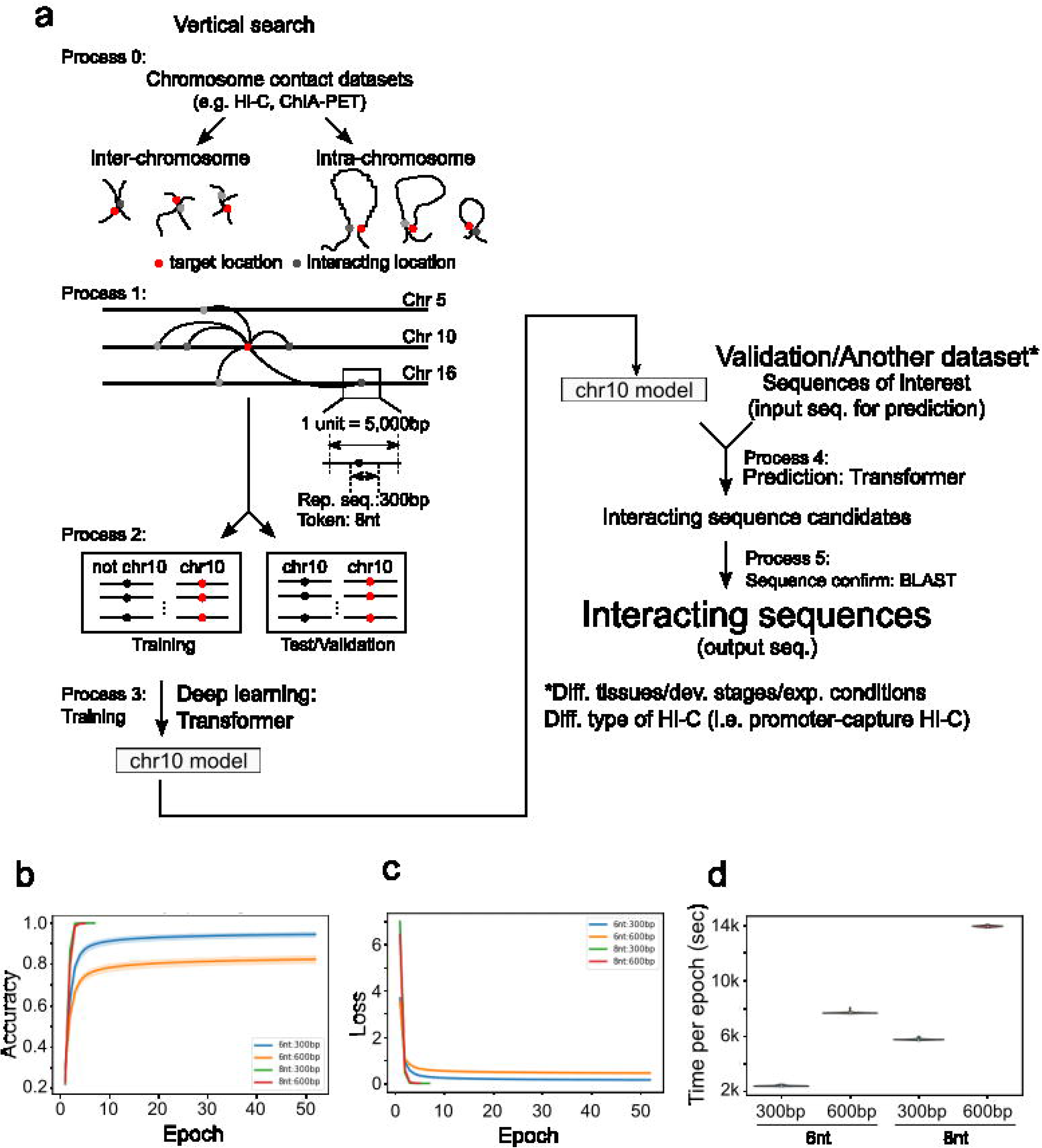
Schematic illustration of a horizontal search for digital Hi-C assay. (a) Schematic illustration of a horizontal search for digital Hi-C assay: process 0: Digital Hi-C assay only requires paired sequence such as chromosome conformation capture basis assay (e.g., Hi-C, ChIA-PET). Inter- and intra-chromosome contacts would be treated separately. Process 1: The sequence mapped to the genome, the mapped location bin in 5,000bp regions and a single 300bp-length representative sequence is selected from the 5,000bp region. The token length is set as eight. Process 2: Apply custom one-chromosome- leave-out rule (i.e., chromosome 10 in this case) to create two datasets; the test dataset contains the paired sequences of inter-interactions with chromosome 10 while the validation dataset contains the paired sequences of intra-interactions on chromosome 10. Process 3: Apply the transformer algorithm to train a model by using the training dataset. Process 4: Predict target sequences (e.g., interacting sequence to the input sequence) by using input sequences (i.e., validation dataset or the sequences of user’s interest) with the trained model in process 3. Process 5: Map target sequences on the genome and confirm the genomic distance between input and target sequences. If the target sequence is mapped onto the genome and its location is less than 1,000,000bp from that of the input sequence, then the target sequences would intersect with the input sequence. (Default: max close distance = 1,000,000bp in human genome) (b, c) The transitions of (b) accuracy and (c) loss up to the 50 epochs in the training process. Two factors were examined: the length of the token (six or eight nucleotides) and representative sequences (300 or 600 base pairs). Create 10 test-training datasets, each of which contains 1,000,000 interactions selected randomly from the original training dataset (see material and methods) and plot the average scores of accuracy and loss on the test- training datasets in the training process. A cut-off threshold to terminate the training process is 0.99 in accuracy. (d) Training time length for each epoch. Y-axis shows the time in seconds (s) to complete a training model for each epoch. Two factors were examined: the length of Token (six or eight nucleotides) and representative sequences (300 or 600 base pairs).

Two interaction datasets; training and validation datasets, were generated from each raw Hi-C dataset followed by the custom one-chromosome-leave-out rule (figure 1a process 2). The removed chromosome is considered as the target chromosome, then the training dataset contained inter-chromosome contacts to the target chromosome whereas the validation dataset contained intra-chromosome contacts on the target chromosome. These interaction datasets contained the paired sequences of genomic loci separated by a comma. Note that the interaction information can be considered separated information per locus; theoretically, these datasets can be split into multiple ways depending on the user’s will (e.g., loci on p/q-arms). A transformer algorithm is applied to create a model per chromosome and dataset (e.g., tissue-/condition-specificity). By the rule of thumb, the training process would be completed by the epoch = 30. The trained model is used for predicting the target sequence (e.g., enhancer) by using the input sequence (e.g., promoter). In the validation process, two criteria must be satisfied to call the prediction successful among the target sequences; the first 50bp-length nucleotides in the target sequence must be a perfect match on the genomic sequence, and it must locate within the fixed-length bp (default 1,000,000bp in human genome case) from the input sequence. Otherwise, those sequences are not called successful (i.e., fail predictions, the locus does not interact with the input sequence). This search approach is called the “vertical search approach”.

The computational code is written in python which is available on GitHub (link). Two notices are mentioned; (1) it takes more than 24 hours to complete a model with “spec of GPUs”, and (2) depending on the GPU environment user used, you may be required to modify the size of chunk data for training depending on the memory capability.

### Horizontal search approach to predict genomic contexts from the result of vertical search approach

A one-dimension (1D) weighted matrix (WM) with 2N-length (default N=5 unless stated) represented the neighbouring regions of a target locus (i.e., the position of target locus = the fifth position in the WM).

In the predictions of enhancer-promoter interactions (EPIs) and CTCF-enriched regions, the WM contains ’1’ in all elements. If the locus has a predicted interaction with the input sequence in vertical search results, multiply the value at the position of the WM with ’1’. Otherwise, multiply the value with ’0’. For example, if all loci in neighbouring regions do not have predicted interactions with the input sequence, the WM should contain 10 zeros. In addition to the neighbouring information, the following two factors are considered: the lowess-normalized GC percentage and the binary information of gene body or intergenic region at the target locus. Those are input data for the random forest (RF) algorithm prediction (in python random-forest package) for genomic context. As validation data for the RF, whether the target locus has an interaction (i.e., EPI information) with the input sequence has been checked for a positive-validation score in the reference dataset.

Balancing the positive/negative ratio in the data, a randomly-selected locus which was not conflicted with reference data is additionally prepared as a negative-validation score. A total of 13 data points (i.e., 12 for input data and 1 for validation scores) were used for the RF. We collected those data at 200 loci which were randomly selected on the target chromosome. After removing duplicated information, the information was shuffled, 90% of the data was used for training to create an RF model and the remained 10% of the data was used for the RF-model performance validation. If training and validation data are different conditions (i.e., different cell lines), then all training data of a cell type was used for the RF to create an RF model and the RF-model performance was measured on the validation data of different cell types. Repeat this process 100 times to compute the performance stats of the following five factors: accuracy, specificity, precision, sensitivity, and F1-score.

In the predictions of TAD-boundary regions, the same strategy is applied with two modified factors. The WM contained the highest values in the middle positions, gradually declined the values at each position toward the edge, and the lowest values in the edge positions. Only WM and lowess-normalized GC percentage at the target locus were considered for prediction.

### GC % normalization per region for prediction of genomic contexts

GC% were computed per unit (i.e. default = 5,000bp length). The GC% within the target region (target loci +/- 1Mbp) are normalized by applying LOWESS (in python lowess package) with option bandwidth=0.4 and polynomial Degree=1.

### Predicting the interactions in promoter-captured Hi-C assay

Promoter-capture Hi-C (PCHi-C) data^19^ was used and contained two types of datasets: PP and PO data in a total of 27 human cell/tissue types. The ‘PP’ data contained ‘promoter-promoter’ interactions whereas the ‘PO’ contain “promoter-other” interactions.

Here, all of the ‘other’ regions in ‘PO’ interactions were assumed as enhancer regions; thus, PO interaction was considered as ‘enhancer-promoter interactions’ Their genomic loci data which have defined interactions were converted into the genomic loci information and bin the interaction positions by 5,000bp. In PCHi-C data, only interactions of genomic loci apart 1Mbp or less were considered. Apply horizontal search to predict PCHi-C interaction [materials and methods]. Since PCHi-C predicted regions (20,000bp on average) were relatively wider than ours (5,000bp), thus, if our results were predicted within the PCHi-C regions, then the prediction was successful. Repeat this process 100 times to compute the average statistical scores; Accuracy, Specificity, Precision, Sensitivity, and F1-score.

### Predicting the CTCF-enriched regions

The data of CTCF-enriched regions are collected from Khoury et al., (2020)^47^. The narrow peak of the control condition was used as validation data. A horizontal search approach is applied to create a model for the CTCF-enriched regions with the input of neighbouring predicting information, lowess-normalized GC% and the status of gene-body/intergenic region. The average length of defined enriched regions is approximately 300bp. If our results (minimum 5,000bp) contained the CTCF-enriched regions, then the prediction was successful. Repeat this process 100 times to compute the average statistical scores; Accuracy, Specificity, Precision, Sensitivity, and F1-score.

### Predicting the TAD-boundary regions

The original data of TAD regions are collected from Rao et al., (2014)^20^. The TAD information is used in the regions defined in the 3D Genome Browser at Northwestern University^48^. Thus, the TAD boundary information is the regions which are not covered by the TAD information and are used for reference data for horizontal search. Custom TAD scores were computed with five neighbouring regions of the target locus, and normalized scores by LOWESS were used as input data for the random forest algorithm together with lowess-normalised GC%. Repeat this process 100 times to compute the average statistical scores; accuracy, specificity, precision, sensitivity, and F1-score.

### Motif search in TAD-boundary candidates

The energy method, so-called BEEML^50^, was employed for identifying the enrichment of transcription factor binding sites (TFBS) followed by previous studies^51^. Briefly, all TFBS information, position weight matrix (PWM), is collected from CIS-BP^52^ and Uniprobe^53^. Due to computing penalty score in the BEEML, the length of PWMs trimmed on both edge sides if the edge of PWM is not conserved well (i.e., max score at a position in PWM is less than 0.5). BEEML score >= 0.09 is considered motif candidates. The regions categorised as TAD-boundary (e.g., N-bp x 5,000bp region) are used for motif search. The occurrence/frequency of PWMs on the target regions is normalized by the length of regions. The enrichment of TFBS is determined by comparing those in the negative datasets that contain the randomly selected 50,000 sequences of 5,000 bp-length from the target genome. Subcategoriezed TAD boundary regions are computed in the same approach per sub-group instead of the whole group.

## Result

### Flowchart of this study

Here, we introduced a digital Hi-C assay by demonstrating the predictions of (1) the long-distance interactions of genomic loci (e.g., enhancer-promoter interactions), (2) the location of CTCF-enriched regions, and (3) TAD boundary regions. In this computational assay, we aimed to create machine-/deep-learning-based models per chromosome by using the genomic-loci interactions data only (i.e., Hi-C dataset).

We achieved this goal by considering predicting the long-distance interactions of genomic loci was conceptually the same as language translating; a sentence (i.e., input sequence) in one language translated to another sentence in another language which was the paired target sequence. We maximized the opportunity for better prediction by considering the following several points. Firstly, we did not employ pre-defined genomic features such as defined enhancer and promoter regions. Instead, we used all raw interactions captured by Hi-C assays (figure 1a process 0). Secondly, we employed shorter-length sequences for input data (e.g., a few hundred base pairs (bp)), each of which was a representative of a wider genomic locus (e.g., a few thousand bp). In particular, we fixed the lengths of input and target sequences to be 300bp which was the represented sequence of a 5,000bp-length genomic locus (discussed below in figure 1a process 1). Thus, the interaction information of our training/validation datasets ought to contain pairs of 300bp- length sequences. We predicted the long-distance interactions of genomic loci by predicting only hundred-bp sequences that were associated with target genomic loci. Thirdly, we applied the customized one-chromosome-leave-out rule (material and methods, figure 1a process 2) for preparing training/validation datasets to predict interactions. We selected a target chromosome, say chromosome 10. The training dataset contained inter-chromosome interactions to the leave-out chromosome 10 whereas the validation dataset contained intra-chromosome interactions onto the removed chromosome. The benefit of this approach was that users could select the sizes of target ‘leave-out’ regions by splitting into multiple sub- groups without losing information (i.e., the range from an entire single chromosome to a single locus). Fourthly, we adopted a sequence-based approach with a deep-learning method, the transformer algorithm^44^ (figure 1a process 3). A sequence-based approach could predict the interactions of genomic loci from a simple and limited number of datasets. The transformer algorithm^44^ was used in natural language processing (NLP). Unlike another popular recurrent neural network such as a Long Short-Term Memory^45^ (LSTM), the transformer algorithm contained an attention process that assessed nucleotide positions, patterns, order, and other unknown sequence features and contexts in given sequences. In the training and validation process, we trained computational models on created training datasets. We utilized the trained models to predict target sequences (figure 1a process 4). As an advantage of the transformer algorithm, the trained model could apply to other datasets even under different conditions such as tissues, developmental stages, different types of Hi-C methods (e.g., in situ or promoter-capture Hi-C) or integrating other high-throughput sequencing datasets such as ATAC-seq, ChIP-seq, GWAS etc, and the sequences of the user’s interest (e.g. input: active promoter region, target: enhancer region) (figure 1a process 5).

Our predicted outputs did not contain any information on genomic features and context because our approach did not use pre-defined information for input data. We addressed this issue by analysing the neighbouring predicted patterns with the GC% in the regions by applying a random forest algorithm^54^ (discussed later). We called this process horizontal search. We used the publicly available datasets of promoter-capture Hi-C, ChIP-seq, and Hi-C assay for training and demonstrated that our digital assay could predict the long-distance interactions of genomic loci, CTCF-enriched regions and TAD-boundary regions efficiently.

In this study, there were different categories among the interactions; hereafter, we categorized the types of interactions in the following manner. A ’captured’ interaction was the one experimentally captured in the Hi-C assay, and a ’predicted’ interaction was the one predicted by our method. If an interaction satisfied the above two categories, we called it a ‘detected’ interaction.

### Suitable conditions for the length and its token length of input sequences

We reconsidered the conditions of the input sequences for the transformer algorithm, especially following two lengths; an input/target sequence and a token (i.e., the minimum length of nucleotides to convert the nucleotides to a numerical index). Previous sequence-based approaches used the same paired information of enhancer and promoter sequences defined in a pioneer study^20, 33, 55^. The majority of those sequences in those datasets overlaid other sequences, thus, influencing the training performance and the prediction results unconsciously and significantly. A previous study reported that the prediction performance declined drastically if such conflicted parts were removed^43^. As a second factor, we wondered about the token length being fixed at six nucleotides (i.e., 6nt) which were used for the predictions with the LSMT algorithm due to computational performances and costs.

However, it was unknown whether the length was optimized for the sequence prediction with the transformer algorithm or our assay. We evaluated the optimization of each condition in terms of prediction accuracy, loss, and computational cost.

Firstly, we optimized the length of input sequences under the fixed-token-length condition. As a model case, we used datasets for chromosomes 10 and 16 in four datasets (HIC001, HIC002, HIC003, and HIC017) from Rao et al., (2014)^20^ (i.e., eight datasets in total). To begin with, we tested the standard conditions; the input-sequence length was 3,000bp with the 6nt-length token. We found that our assay took more than 24 hours to generate/update a model per epoch, resulting in those standard conditions were not realistic for our approach (data not shown). Next, we varied the lengths of input sequences as follows; 100, 200, 300, 600, and 1,200bp. We found that the maximum accuracy declined as the length of the input sequence become longer (supplemental figure 1a, g). For instance, on chromosome 10, we found that the maximum accuracy at 20 epochs reached 0.85, 0.7, 0.6, and 0.55 for 100bp, 200bp, 300bp and 600bp, respectively. As correlated with those trends, the longer input sequence returned larger scores in loss (supplemental figure 1b, h).

Additionally, we found that the length of the input sequence correlated with the time length of computations exponentially (supplemental figure 1c, i). We could find similar trends at token length was 8nt (supplemental figure 1d-f, j-l). As considered less redundancy in input sequences and avoided mammalian ‘super-conserved region’ which might hamper our predictions, the input sequence length ought to be at least 300bp.

Next, we evaluated the effect of the token length on the prediction performance under the 300bp-length input sequence. We varied the length of a token as follows; 4, 6, 8, and 9nt. We again evaluated the training performance by the same three factors.

Unexpectedly, we found that the token length of 8nt gave the highest accuracy; the longer token the better prediction. At nine or longer nucleotides (nt) in length, however, it took more than 24 hours per epoch to learn. We noted that accuracy and loss for the learning process could reach accuracy > 0.98 and loss < 0.05 under 300bp-length input sequence and 8nt- length token on chromosomes 10 and 20 (supplemental figure 1m-n, p-q). The computational cost also increased exponentially as the length of tokens was longer on both chromosomes 10 and 20 (supplemental figure 1o, r). We observed similar trends in the case of the 600bp-length input sequence (data not shown).

The combination of the ‘local’ best conditions might not be suitable condition overall. Hence, we directly compared the prediction performance with the combinational conditions; 300bp vs 600bp in input sequence length and 6nt vs 8nt in token length. As a result, we found that the highest accuracies and lowest loss returned at 8nt-length tokens with 300bp and 600bp-length input sequences and terminated their computational training earlier than other conditions because it reached saturation conditions (figure 1b, c). By considering computational costs, we found the shorter sequences were, the quicker computation was completed per epoch (figure 1d). Thus, we concluded that the appropriate lengths of the input sequence and token for our transformer-basis assay were 300bp and 8nt, respectively.

### Comparison of prediction and captured interactions (chromosomes 5, 10, and 16 in GM12878 cell lines of in situ Hi-C)

We evaluated the prediction performance of our digital assay by demonstrating how efficiently our trained models could predict the captured interactions in the Hi-C assay. To conduct this, we used the HIC-001 dataset of GM12878 cells in Rao et al., (2014)^20^ and here used chromosomes 5, 10 and 16 as model cases. As we mentioned above, our training datasets contained inter-chromosome interactions whereas the validation datasets contained intra-chromosome interactions (figure 1a process 2, materials and methods). Each dataset contained the paired 300bp-length sequences which represented input and target sequences. We randomly selected 200 loci (i.e., input sequences for prediction) on each of the chromosomes for validation performance stats. We computed the performance stats in the following five factors; accuracy, precision, recall, sensitivity, and F-1 scores, by counting the number of predicted and captured interactions of the target loci (materials and methods). We simplified to visualizing the predicted/reference interactions with a ‘bar’ on the locus if the target locus (i.e., target sequence) predictably interacted with the input locus (i.e., input sequences) (figure 2a).

**Figure 2:**
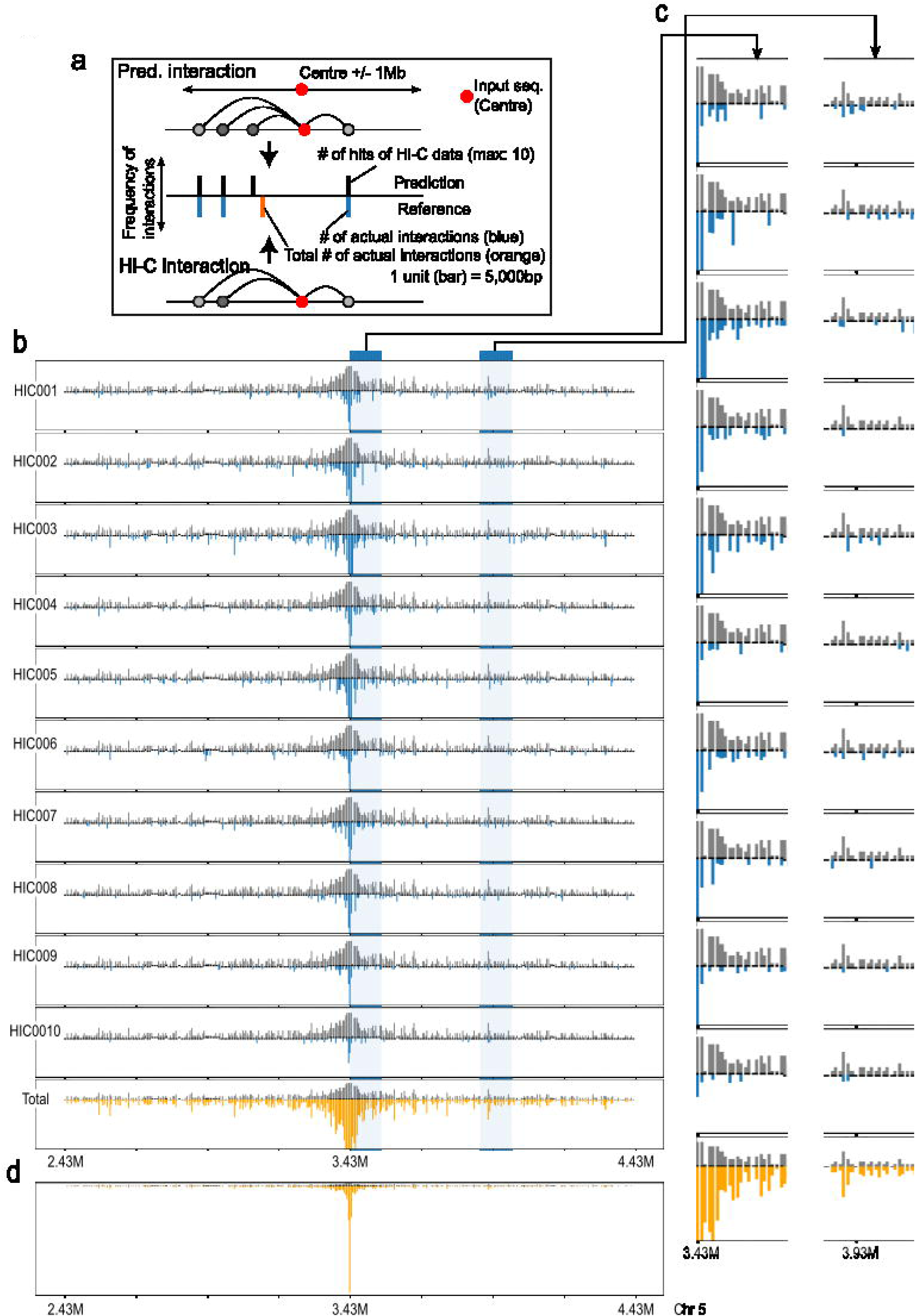
Visual comparison between predicting and reference interactions of genomic loci. **(a)** Illustration of visualizing long-distance interaction of genomic loci. When the locus has interactions with the input sequence (the centre position of each bar figure), the locus has a bar (black-colour bar for predicted interactions on the top part whereas blue-colour bar for reference interactions on the bottom part). The heights of predicted interactions indicate the number of experimental assays having the interactions (max: 10). The height of reference interactions indicates the Knight-Ruiz (KR)-normalized score^49^ of the interactions between input and target loci. (b) A representative result of predicted interactions over 10 different reference datasets (HIC001 - HIC010 from Rao et al., 2014^20^) in the range of 2,430,000bp to 4,430,000bp on chromosome 5. The model trained on the training dataset of HIC001 is used for prediction on each of the listed HIC datasets. The input sequence locates in 3,430,000bp on chromosome 5. Each row figure contains the comparative results between predicted and reference interactions. The name of reference data (e.g., HIC001) is shown in the left position of each bar figure. The “total” (or orange-colour bar) in the last column indicates the sum of 10 reference scores above (HIC001 - HIC010). (c) Two zoom-in areas (white-blue areas) in the representative result (b). The order of rows should be identical to those in (b). (d) The zoom-out version of ‘total’’ result in (b).

Firstly, we looked at predicted and captured interactions around the region of 3.43Mb +/- 1.0Mb on chromosome 5 as a represented result and visualized the 10 results per reference data (figure 2b, HIC001 - HIC010). We found that predicted results (i.e., the grey- colour bar on top) could completely detect the raw interactions (i.e., the blue-colour bar on the bottom) within the target regions (figure 2c (single data with HIC001), supplemental figure 3c). Whereas those results also showed that we predicted a lot more interactions than those of captured interactions in each reference data (figure 2b-c, e.g., a predicted vs HIC001 top blue bar). Consistently, this result was shown in poor performance stats; 0.4 in accuracy, 0.4 in F-1 score (figure 3b). This trend was not difference in different datasets in the same cell lines (e.g., HIC002), at different loci (data not shown), and different chromosomes (supplemental figure 2a-c,d-f). It is known that single Hi-C experiments could limitedly capture the number of true interactions; indeed, they might detect only 10-15% of genuine interactions^20^. To confirm this idea, we made a dataset of ‘total’ interactions (i.e., orange-colour bar) which contained all captured interactions of the targeted 10 reference datasets (i.e., HIC001 – HIC0010) for the target region. We labelled the results with ‘total’ instead of a specific reference dataset name. We found that our assay efficiently predicted the total interactions of the target locus on this dataset (figure 2b, c bottom); the performance stats on total reference data showed above 0.8 scores in all categories (figure 3b).

**Figure 3:**
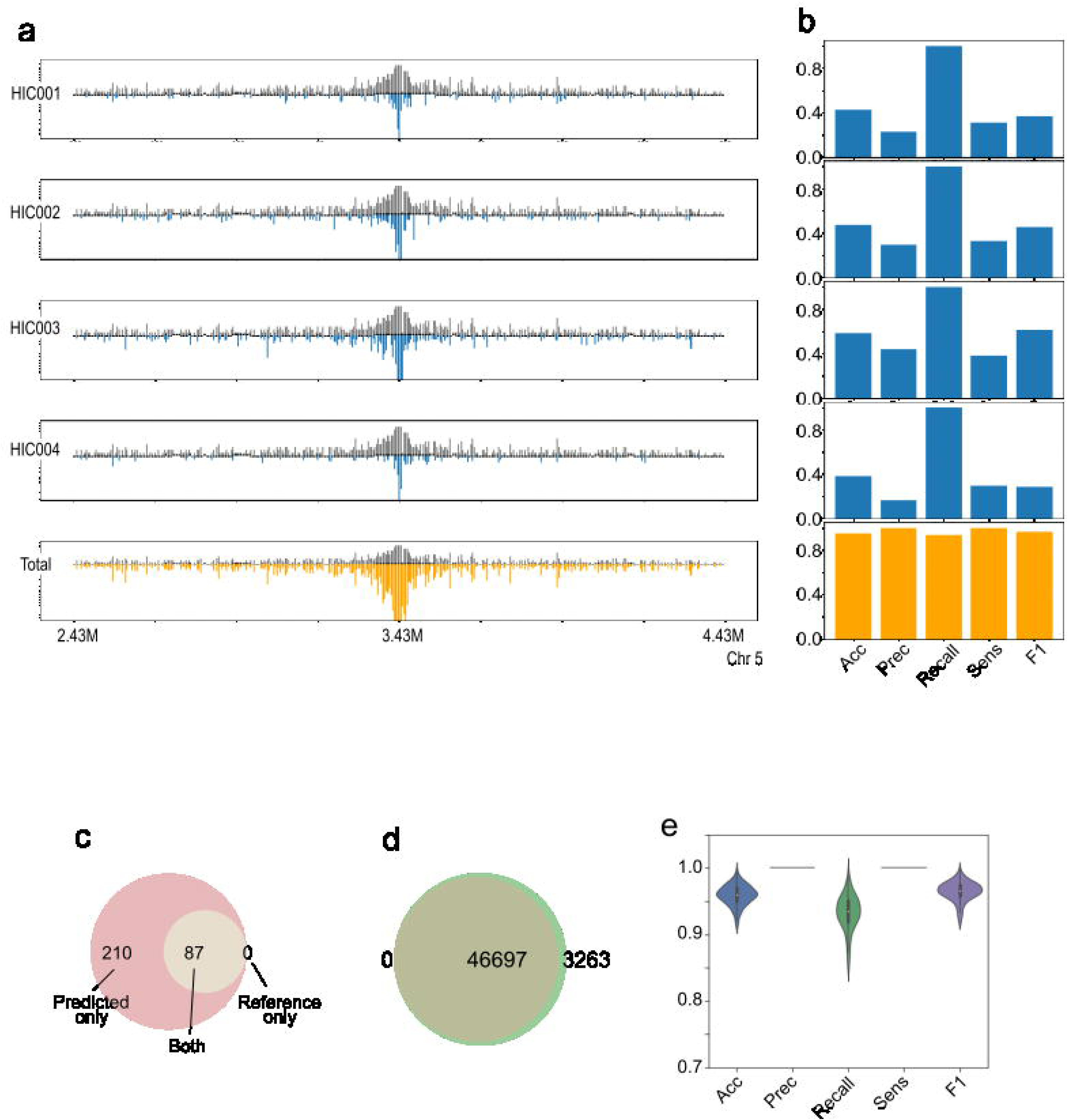
Performance statistics of prediction on chromosome 5. (a) A representative stats performance results of predicted interactions with 4 different reference datasets only (HIC001 - HIC004 from Rao et al., 2014^20^) in the range of 2,430,000bp to 4,430,000bp on chromosome 5. The “total” (or orange-colour bar) in the last column still indicates the sum of 10 reference scores above (HIC001 - HIC010). (b) Prediction performance stats between predicted (positive/negative) and reference (true/false) interactions for each dataset. The input sequence locates in 3,430,000bp on chromosome 5. The performance stats are measured following five factors; accuracy, precision, recall, sensitivity, and F-1 score. Each row result corresponds to the reference dataset in the same row in (a) (c) Venn diagram of the predicted and reference interactions for the target input sequence at 3,430,000bp. Predicted and reference interactions whose locations are less than 1Mb from that of the input sequence are considered. In (b), reference data of HIC001 is considered. In (c), reference data of ten (HIC001 - HIC010) are considered. (d) Venn diagram of the predicted and reference interactions at the 200 loci. The 200 loci are randomly selected on the target chromosome. Reference data of ten (HIC001 - HIC010) are considered. (e) Overall prediction performance stats between predicted (positive/negative) and reference (true/false) interactions at the 200 loci.

Furthermore, we looked at all selected 200 loci and found the result still showed our assay predicted 93% of reference interactions (figure 3d); consistently, it supported with high-performance stats; at least 0.95 scores in all categories (figure 3e). Although we closely looked at the prediction patterns to investigate any correlated or biases of the interaction positions that existed between predicted and captured interactions, we could not find particular correlations between the two types of interactions. Taken together, we concluded that our method could efficiently predict the captured interactions in the Hi-C assay; especially those redundantly captured by multiple assays.

### Efficiently predicting the interactions of genomic loci in datasets by using the models trained on different datasets

In practical issues, it would be desired to predict long-distance interactions of genomic loci even if Hi-C datasets were not fully available for particular conditions of interest. Next, thus, we aimed to evaluate how efficiently a model trained on a dataset could predict the long-distance interactions of genomic loci in different datasets (e.g., different cell lines, experimental conditions etc).

We first address this by applying the HIC001-trained models, as a model case, to predict long-distance interactions in the NHEK-cell-line datasets (i.e., HIC065 - HIC068). As a represented result, we showed all predicted and reference interactions around 42.56Mb +/- 1Mb on chromosome 5 (figure 4a). Around the target regions, the prediction performance stats had > 0.6 scores in both accuracy and F1-score, indicating that the HIC001-(GM12878- cell-line) trained model was a good prediction model for the long-distance interactions in NHEK-cell-line (figure 4b). To be consistent with this result, we showed that regions at 8.435Mb +/- 1Mb on chromosome 16 had similar trends in their prediction results (figure 4c) and their overall performance stats remained above 0.6 in the F-1 score (figure 4d). We looked at all randomly selected 200 loci on chromosomes 5 and 16. We found the result showed a relatively lower prediction rate (38.0% for chromosome 5 and 30.2% for chromosome 16) (figure 4e, f). The prediction performance stats on chromosome 5 showed 0.6 scores in F-1 score (figure 4g) whereas that on chromosome 16 was around 0.5 in F-1 score (figure 4h). The overall trends on chromosome 10 were similar to those of chromosome 5 (supplemental figure 3a,b). A represented result around 33.295Mb +/- 1Mb on chromosome 10 and their performance stats around the target regions were described in supplemental figure c.

**Figure 4:**
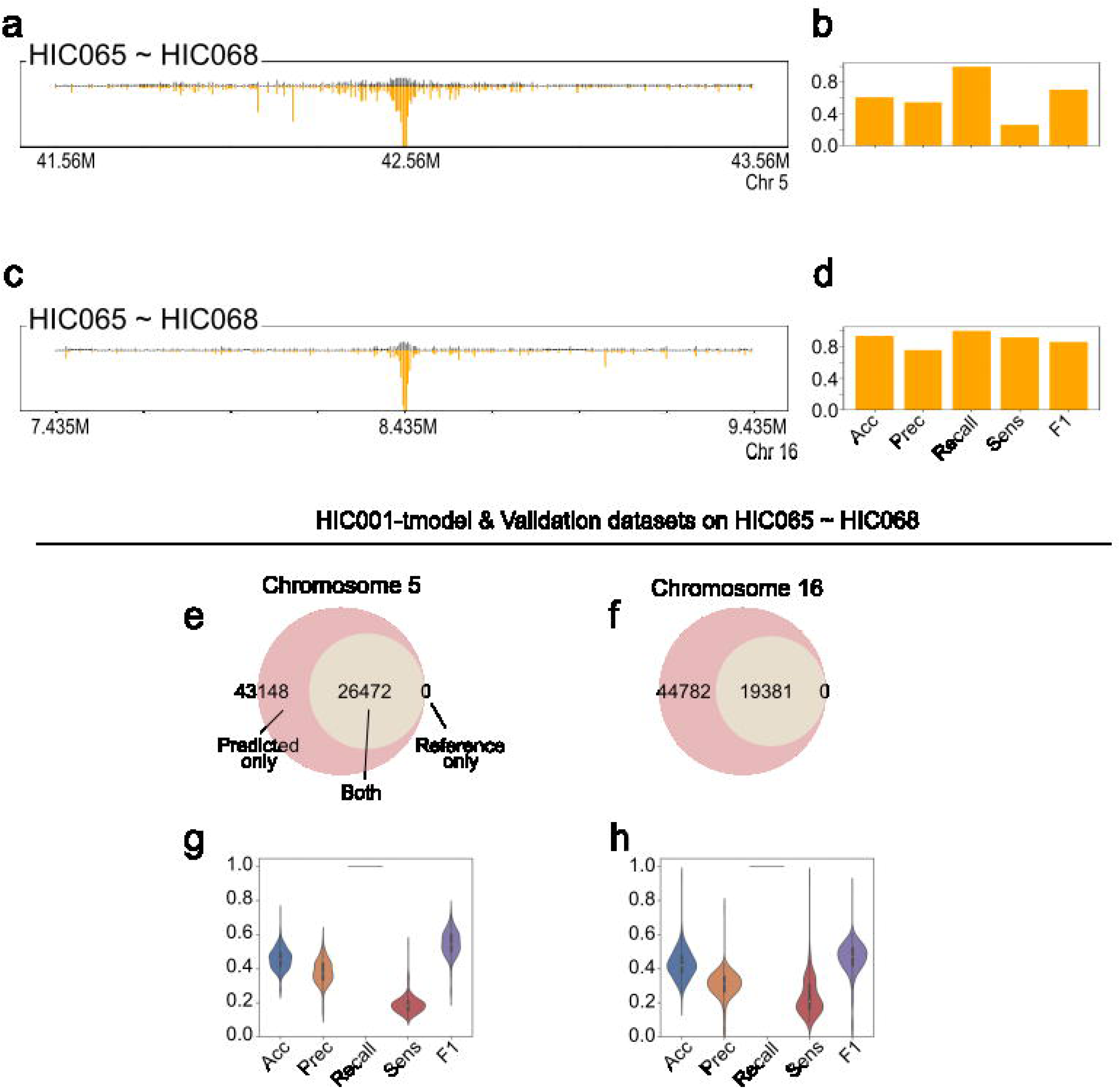
Efficiently predicting the interactions of genomic loci in datasets by using trained models in different datasets (on different chromosomes 5 and 16 with HIC65). (a-d). A representative stats performance results of predicted interactions with four different reference datasets only (a-b: HIC065 - HIC068 from Rao et al., 2014^20^) in the range of 41,560,000bp to 43,560,000bp on chromosome 5. The input sequence locates in 42,560,000bp on the chromosome. (c-d: HIC065 - HIC068) in the range of 7,435,000 bp to 9,435,000 bp on chromosome 16. Prediction performance stats between predicted (positive/negative) and reference (true/false) interactions for each dataset. The performance stats are measured following five factors; accuracy, precision, recall, sensitivity, and F-1 score. (e, f) Venn diagram of the predicted and reference interactions at the 200 loci on chromosomes (e) 5 and (f) 16. The 200 loci are randomly selected on the target chromosome. The model is trained on HIC001. The trained model is used to verify the prediction performance on four reference data (HIC065 - HIC068). (g, h) Overall prediction performance stats between predicted (positive/negative) and reference (true/false) interactions at the 200 loci. The performance stats are measured following five factors; accuracy, precision, recall, sensitivity, and F-1 score.

Those prediction performances were worse than those in previous sections (e.g., training and validation data have been used for the interactions in the same cell line). We investigated the causative factors which returned poorer prediction performance. Hence, we hypothesized these cases might be the outcomes in which the number of experimental datasets was limited (e.g., there were four datasets for NHEK-cell-line datasets while the GM12878 we tested above had 10 datasets.), thus, could not experimentally recapture total interactions of genuine positives. We verified the hypothesis by using the GM12878-cell-line datasets and creating two sets of four reference datasets from the GM12878-cell-line datasets; the first set contained the HIC002 - HIC005 datasets, and the second set contained the HIC006 - HIC009 datasets. Although we applied the same HIC001-trained model which showed excellent prediction performance in the previous section (figure 2), we found that the prediction performances on both data groups declined to a similar level as those on NHEK-cell-line datasets (supplemental figure 4a-e). We found that lower prediction rates (50.6% for HIC002 - HIC005 (supplemental figure 4a) and 33.3% for HIC006 - HIC009 (supplemental figure 4b)) while lower prediction performance stats were on 0.5 - 0.7 in F1 scores (supplemental figure 4c, d). We noted that our model still predicted long-distance interactions of genomic loci (supplemental figure 4e), and the performance stats (e.g., accuracy and sensitivity etc) were lower in comparison to those having a large number of reference datasets. Those results indicated that the number of available reference datasets might significantly influence the systematic scorings for our prediction performances on long-distance interactions.

To support this hypothesis further, we verified the prediction performances of HIC001-trained models on the three additional datasets: IMR90, HMEC, and HUVEC-cell-lines^20^. We noted that these datasets contained in the range of four to eight datasets. As a model case, we focus on chromosome 10. We found that our HIC001-trained models could predict all captured interactions at randomly-selected 200 loci in those three cases (supplemental figure 5a,b,d), consistent with those of NHEK-cell-line case (supplemental figure 5c). This time, we found that prediction performance based on the F1-score was slightly lower than those of the HIC065s datasets, but its F1-score remained above 0.6 under those conditions (supplemental figure 5e-h). The represented results for each dataset were shown in supplemental figure 5i-l. Taken together, we showed that our approach allowed the models trained on a dataset to apply to predict long-distance interactions on different datasets efficiently. Due to lack the number of reference datasets, the standard prediction performance might not be successful; however, all results suggested that our strategy and digital assay might predict genuine interactions of genomic loci even though the Hi-C datasets in the cell lines might not be available in specific conditions.

**Figure 5:**
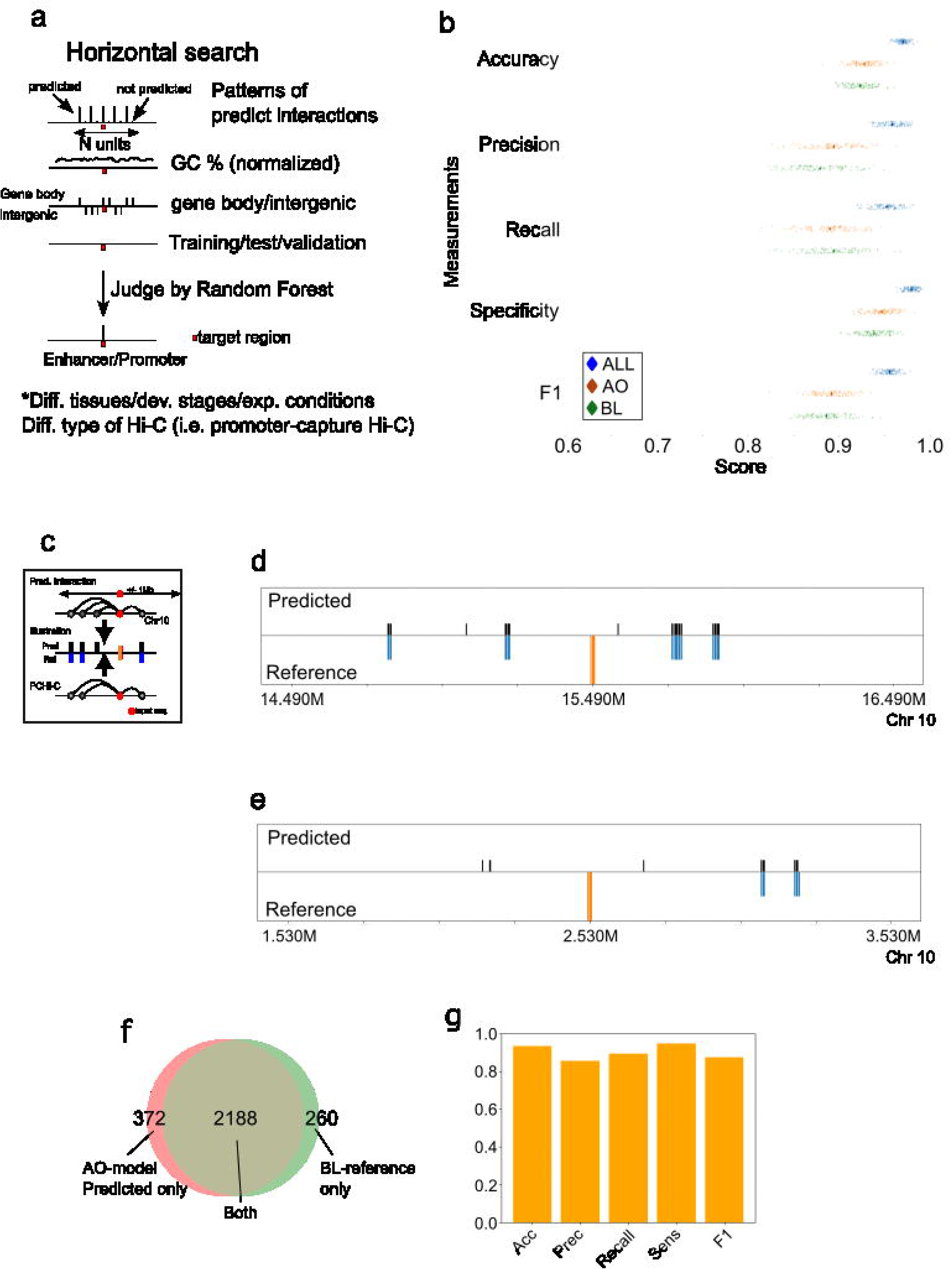
Predicting enhancer-promoter interactions (EPIs) by horizontal search approach. (a) Schematic illustration of a horizontal search for predicting enhancer-promoter interactions. Predicted interactions around a target locus are considered ten input data (N=10). Normalized GC % in the target locus (total 10 data points) and the information on whether the locus is in a gene body or not. A total of 21 data points is used to create a model with a random forest algorithm and predict whether the target locus is categorized as genomic context (e.g. promoter, enhancer). the datasets of promoter-captured Hi-C (PCHi-C) [Jung et al., 2019] are used as reference datasets. (b) Plots for prediction performance statistics for EPIs in PCHi-C. The performance stats on three datasets; ‘ALL’, ‘AO’, and ‘BL’, are shown as represented results. ‘ALL’ contained all interactions across 27 tissues in PCHi-C datasets. 90% of the data are used for training and the random-forest model is tested on the remained 10% of the data. Shuffle the data and repeat the performance validation process 100 times to compute performance stats of five factors; accuracy, precision, recall, sensitivity, and F-1 score. (c) Illustration of visualising the predicted and reference of enhancer-promoter interactions. An input sequence is marked orange-colour bar. If the locus interacts with the input interactions, then the locus would have a coloured bar (black colour for predicted interaction and blue colour for reference interaction.) Only loci within 1,000,000bp from input sequences are shown. (d) Representative result of enhancer-promoter interactions in the range of 14,490,000bp to 16,490,000bp on chromosome 10. The location of the input sequence is 15,490,000bp on the chromosome. The model was trained on 90% of the AO data and validated the prediction performance of the model on the remained 10% of the AO data. (e) Representative result of enhancer-promoter interactions in the range of 1,530,000bp to 3,530,000bp on chromosome 10. The location of the input sequence is 2,530,000bp on the chromosome. The model was trained on 100% of the AO data and validated the prediction performance of the model on the remained BL data. (f) Venn diagram for the overlap of predicted and reference data at the 200 loci on chromosome 10. The 200 loci are randomly selected on the target chromosome. The model was created on the AO data above. The created model was used to verify the prediction performance on the BL data. (g) Overall prediction performance stats between AO-model prediction (positive/negative) and BL-reference (true/false) interactions at the 200 loci.

### Toward identifying genomic contexts from our predicted results

In the following sections, we addressed whether our digital assay could predict the genomic contexts and compartments from its prediction results. One of our ultimate goals was to develop a digital assay which allowed us to predict the long-distance interactions of genomic loci even under all high-throughput experimental results (e.g., Hi-C, ATAC-seq, ChIP-seq etc) have not available for the target and specific conditions of interest. Massive lines of evidence showed that genomic context (e.g., enhancer-promoter interactions) and genomic three-dimensional (3D) architecture (e.g., topological associating domains (TAD)) could be detectable from Hi-C data by integrating other high-throughput data such as ATAC-seq and ChIP-seq and applying advanced statistical and computational methods. For example, those genomic compartments could be defined bioinformatically by using chartink patterns of interaction frequency in analysed Hi-C data^9^. This indicated that Hi-C data ought to contain all information. In addition, hypothetically, it is possible to extract such genomic contexts and 3D architectures from Hi-C data with appropriate advanced strategies without using other types of high-throughput data. Up to this point, our digital assay demonstrated that it could flexibly and reliably predict the long-distance interactions of genomic loci by using the paired sequences directly (i.e., raw captured interactions) from Hi-C datasets. In those prediction results, we observed that some genomic loci tended to have ’dense’ interactions in their neighbouring whereas others had ’sparse’ interactions. We speculated that the prediction patterns around a target genomic locus might correlate with specific genomic contexts.

To address this, we introduced an additional function, so-called the horizontal search approach, to predict genomic context (e.g., enhancer-promoter interaction) (figure 5a) [materials and methods]. Although the targeting genomic features were slightly optimized for each study case, the overall strategy in the prediction of these features was the same. Briefly, this horizontal search approach considered 2N-length neighbouring regions (default N = 5) around a target locus (i.e., target locus = the fifth position in the 2N-region). Each position of the 2N region contained the predicted/un-predicted information of vertical search results. In addition to the target local region, we consider two additional reliable factors, normalized GC percentage and binary information of gene body or intergenic region at the target locus. All information was used to create a Random-Forest (RF) model and judge whether the new targeted locus had genomic contexts. In the following sections, we investigated whether our digital assay could predict the genomic contexts and compartments from its prediction results. To conduct this, we applied this horizontal search and categorized whether the target locus had genomic contexts or not. We aimed at the following three contexts: enhancer-promoter interactions, CTCF-enriched regions, and TAD boundary regions.

### Predicting enhancer-promoter interaction by using promoter-captured Hi-C interactions

As a first example to predict genomic contexts, we applied a horizontal search approach to predict enhancer-promoter interaction. In this approach, we considered neighbouring predicted patterns with two additional factors: normalized GC percentage and binary information of gene body or intergenic region at the target locus [materials and methods]. Information from 13 data points in total was used to create an RF model and judge whether the new targeted locus had EPIs at a target locus.

We evaluated whether the horizontal search approach could efficiently predict the enhancer-promoter interactions (EPIs). We used the datasets of promoter-capture Hi-C (PCHi-C)^19^ as reference data. In addition to the 27 original datasets, we created an additional dataset; “ALL’’ contained all EPIs of 27 human tissues. In those datasets, the authors provided two types of interactions of genomic loci: promoter-promoter (‘pp’) and promoter-other (‘po’) interactions. We assumed and used the “po” data representing EPIs. We again used HIC-001-trained models to predict long-distance interactions on chromosome 10 as a model case. We considered 90% of the data for training and remained 10% for validation and computed the performance stats each time [materials and methods]. We repeated this process 100 times to generate the overall performance stats. As represented cases, we showed the performance stats on the following three cases: “AO”, “BL”, and ALL (figure 5b). We mentioned that the “ALL’’ case tended to have higher scores on all performance stats because they contained much larger positive datasets which made the trained model better. Although the performance stats were slightly better on the AO dataset compared to those on the BL datasets, the difference was neglectable. Their F-1 score showed > 0.85, indicating our models-based horizontal search was excellent to capture EPIs. We closely looked at the details of the results in the representative results. We showed the predicted (black-colour bar on top) and reference interactions (blue-colour bar on bottom) around the regions around 15,490,000 bp (input sequence indicating orange-colour bar) on chromosome 10 (figure 5d). We noted the black/blue-colour bar indicated that the locus had predicted/reference interactions to the input sequence (orange-colour bar), respectively (figure 5c). Although the regions with the EPIs in PCHi-C data tended to be wider than ours (e.g., average length of such regions was 25,000 bp), we showed our horizontal search could predict the EPIs regions efficiently at a single locus.

Furthermore, to address generality, we asked whether a trained model in a cell type could predict EPIs in other cell types. We addressed this by demonstrating the RF model of the ‘AO’ could predict EPIs in the ‘BL’ dataset (figure 5e), We showed a representative result of around 2,530,000 bp on chromosome 10. The result showed that a horizontal approach could predict the EPIs in the regions efficiently. We mentioned that this region does not contain the ‘AO’ dataset; thus, it was also supported that a trained model in a cell type could be used for predictions in other cell types. We emphasized that some of the interactions apart more than 750,000 bp between the loci, and our model could precisely predict the interactions within 5,000 bp resolution. We applied this strategy to predict EPIs at 200 loci which were randomly selected on the chromosome. We found that 85.3% of AO-model predicted results could capture EPIs in BL-dataset (figure 5f), and the prediction performance stats showed the model was sufficiently good enough for the prediction (figure 5g). Further, we applied the same AO-model to other cell lines in other datasets of Jung et al., (2019)^19^ (supplemental figure 6a-aa). The prediction rate could vary between 80.5 and 86.7%, which might reflect the variability and similarity of cell-line specific features. These results also showed that a trained model in a cell type could predict to EPIs in other cell types efficiently, indicating that even in the case of fewer numbers and small regions of EPIs, our predictions could predict the EPIs efficiently. Taken together, we concluded that our horizontal search approach could efficiently predict enhancer-promoter interactions. Further, we could show that our approach had general versatility as an application and predict EPIs in the situation of the cell type, not having reference data. Those results suggested that the horizontal search approach could efficiently predict the enhancer-promoter interactions (EPIs), and the trained model on a dataset could be applicable to predict the interactions in different datasets (e.g., different cell lines).

**Figure 6:**
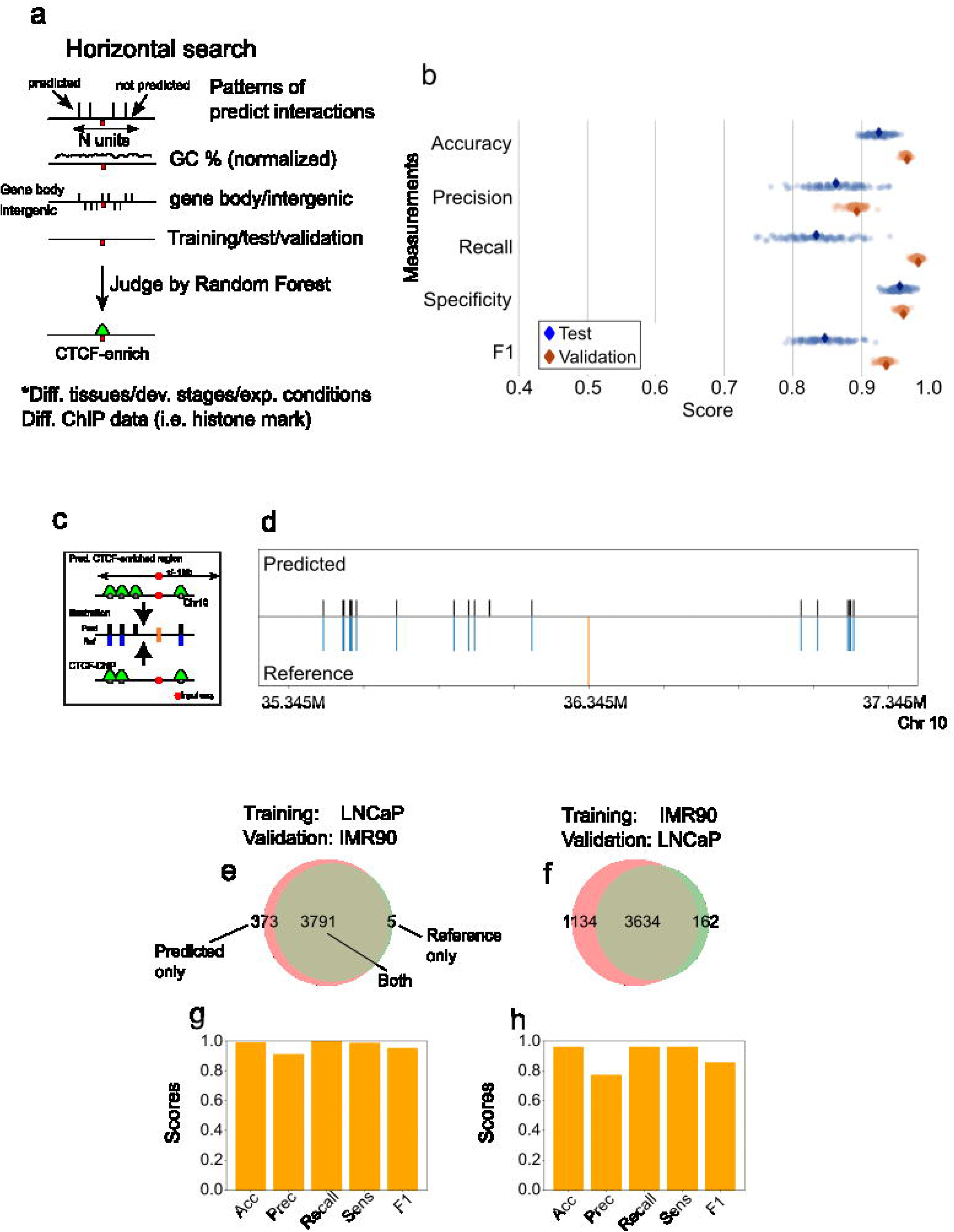
Predicting CTCF-enriched regions by horizontal search approach. (a) Schematic illustration of a horizontal search for predicting CTCF-enriched regions. Predicted interactions around a target locus are considered ten input data (N=10). Normalized GC % in the target locus and the information on whether the locus is in a gene body or not. A total of 12 data points is used to create a model with a random forest algorithm. Use the trained model to predict whether the target locus is categorized as genomic context (e.g., CTCF-enriched regions). The ChIP-seq datasets for CTCF^47^ are used as reference datasets. (b) Plots for prediction performance statistics for CTCF-enriched regions from the ChIP-seq datasets. The performance stats on IMR90 are shown as represented results. 200 loci are selected randomly from the target chromosome. 90% of the data on the chromosome are used for training and the random-forest model is tested on the remained 10% of the data. Shuffle the data and repeat the performance validation process 100 times to compute performance stats of five factors; accuracy, precision, recall, sensitivity, and F-1 score. (c) Illustration of visualising the predicted and reference of CTCF-enriched regions. An input sequence used for predicting long-distance interactions of genomic loci is marked orange- colour bar. If the locus is categorized as CTCF-enriched regions, then the locus would have a coloured bar (black colour for predicted interaction and blue colour for reference interaction.) Only loci within 1Mb from the input sequences are shown. (d) Representative result of enhancer-promoter interactions in the range of 35,345,000 bp to 37,345,000 bp on chromosome 10. The location of the input sequence is 36,345,000 bp on the chromosome. The model was trained on 90% of the IMR90 data and validated the prediction performance of the model on the remained 10% of the IMR90 data. (e, f) Venn diagram for the overlap of predicted and reference data at the 200 loci on chromosome 10. The 200 loci are randomly selected on the target chromosome. The model was created on the training data and validated their performance on validation datasets with the following combinations: (e) the training data are LNCaP cell data, and the validation data are IMR90 cell data while (f) the training data are IMR90 cell data and the validation data are LNCaP cell data. (g, h) Overall prediction performance stats at the 200 loci. (g) has the training data are LNCaP cell data and the validation data are IMR90 cell data while (h) has the training data are IMR90 cell data and the validation data are LNCaP cell data

### The regions having no predicted interactions represent the CTCF-enriched regions with specific prediction patterns

Next, we addressed whether the horizontal search approach could predict another genomic context (figure 6a); CTCF-enriched regions. In our predicted result for long-distance interactions, we found some regions did not contain any prediction. Since Hi-C data could not capture any or a fewer number of interactions of genomic loci around the topological boundary domain (TAD) boundaries, we speculated the regions without predicted interactions in our results might correlate with the TAD boundary regions. It has been known that the boundary regions correlate with the enrichment of the CCCTC binding factor binding sites (CTCF)^9, 56^; additionally, the number of CTCF-enriched regions determined experimentally were larger than those of TAD boundary regions, hence, could be used as a positive control to confirm a prediction performance. Thus, we demonstrated, firstly, the horizontal search approach could efficiently predict CTCF-enriched regions (materials and methods). To conduct this, we used the datasets of ChIP-seq for CTCF^47^ for training and validating our predicted results; the authors provided the datasets of two cell types; IMR90 and LNCaP. We created an RF model by using 90% of the data for training and validated the prediction performance by using the remaining 10% on the CTCF dataset. We evaluated the prediction of the RF model by computing the performance stats of the five measurements. We repeated this process 100 times to compute the performance stats. As a result, we found our models showed 92.9% accuracy, 89.4% precision, and 85.1% F1 on chromosome 5 in LNCaP-cell (figure 6b). Even though we tested this procedure on chromosomes 10 and 16 in the same dataset, these statistical scores remained high (data not shown).

Likewise in the case of PCHi-C prediction, we further demonstrated a broader application by showing those trained models on the LNCaP dataset could efficiently predict the CTCF-enriched regions on another dataset, the IMR90 dataset or vice versa. As a result, we found that the accuracy and other measurable factors remained high; 0.96 in accuracy and 0.94 in F-1 score on chromosome 5. We also confirmed a similar level of predicted performance was conducted on chromosomes 10 and 16. We showed a representative result of around 36,345kbp on chromosome 10 (figure 6d). As we expected from CPHi-C cases, the results showed our prediction detected CTCF-enriched regions in reference data nearly perfectly at this target locus. Further, we applied this strategy to predict CTCF-enriched regions at randomly-selected 200 loci on the chromosome 10. We found the LNCaP-based model could predict 91% success ratio on the dataset of IMR90 (figure 6e,g). Its performance stats showed more than 0.9 scores in all categories. The other chromosomes 5 and 16 also showed the similar trends (supplemental figure 7a,c,e,g). We also validated the other case; predict with IMR90-based model and validate on LNCaP data. Due to smaller size of data in IMR90, the prediction rate was slightly lower (76.2%) on the chromosome 10 (figure 6f,h), and the performance stats showed still 0.8 scores, indicating the model was excellent. The other results on two chromosomes 5 and 16 also showed the similar trends (supplemental figure 7b,d,f,h). Those results suggested that the horizontal search approach could efficiently predict the CTCF-enriched regions. Although the size of positive data might influence the prediction performance, the trained model on a dataset could be applicable to predict the interactions in different datasets.

**Figure 7:**
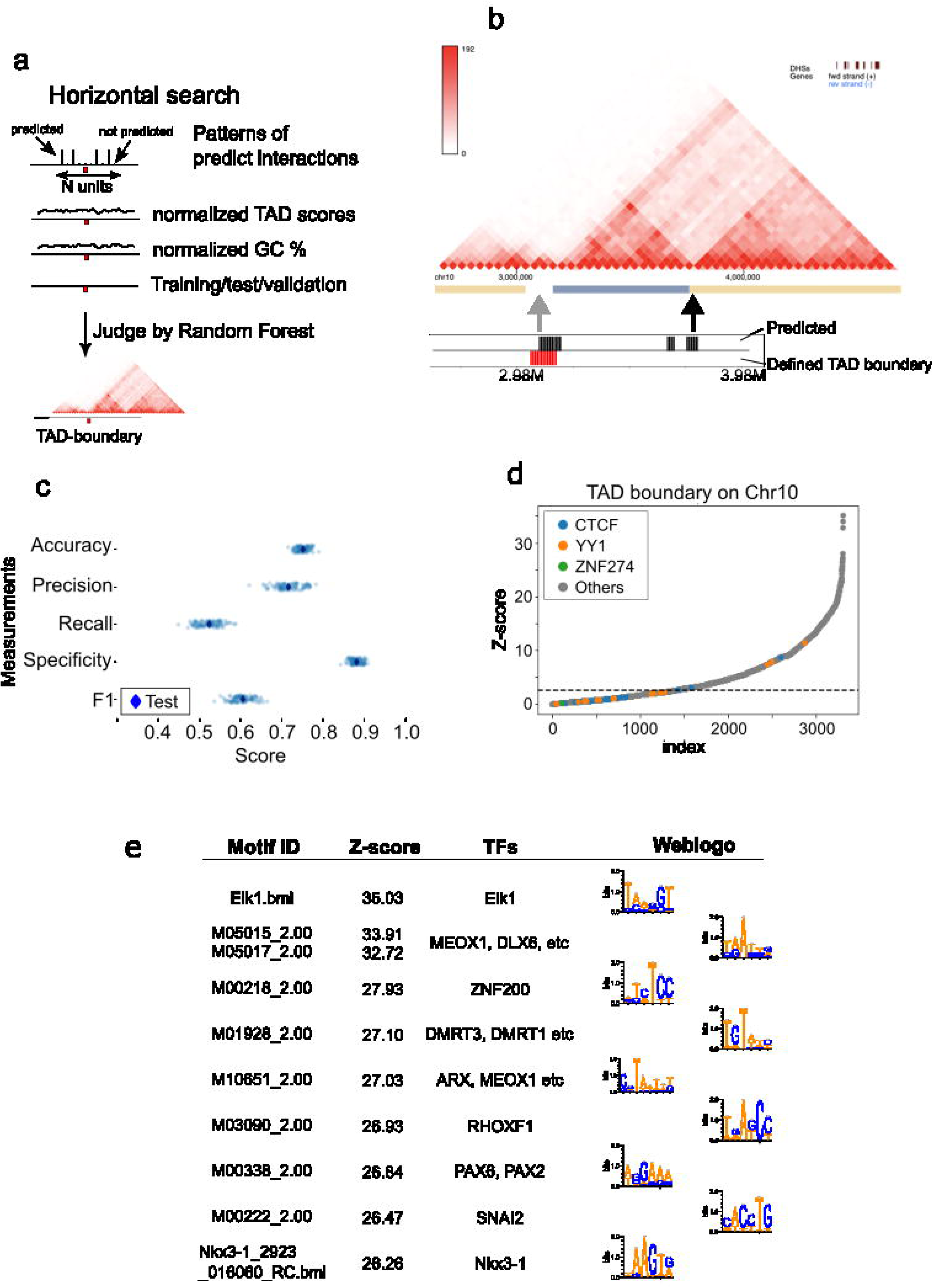
Predicting TAD-boundary regions by horizontal search approach. (a) Schematic illustration of a horizontal search for predicting TAD-boundary regions. Predicted interactions around a target locus are considered ten input data (N=10). Normalized GC % in the target locus is additionally considered. A total of 11 data points is used to create a model with a random forest algorithm. Use the trained model to predict whether the target locus is categorized as genomic context (e.g., TAD-boundary regions). The TAD boundary information as reference data is collected from the 3D Genome Browser at Northwestern University. (b) The location of predicted regions in the range of 2,980,000bp (the location of input sequence) to 3,980,000bp on chromosome 10. In the top image (triangle-shaped image), reference data of TAD and its boundary regions are visualized by using the 3D Genome Browser. The ochre-/blue-colour indicates the defined TAD regions in the original Rao et al., 2014. The ‘white-colour’ gap region between ocher- and blue-colour boxes around 3,000,000bp is defined as the TAD-boundary region in this study. On the bottom, the predicted TAD boundary regions indicate the black-colour bars while reference TAD boundary regions indicate the red-colour bar. (c) Plots for the prediction performance statistics for TAD-boundary regions from the GM19878 in Rao’s datasets. The performance stats on the dataset are shown as represented results. 90% of the data on the target chromosome are used for training and the random-forest model is tested on the remained 10% of the data. Randomly 200 loci are selected from the target chromosome. Shuffle the data of TAD boundary regions and repeat the performance validation process 100 times to compute performance stats of five factors; accuracy, precision, recall, sensitivity, and F-1 score. Each dot represented the individual score of a single model (i.e., single trial). (d) Motif search in TAD boundary regions on chromosome 10. Plot transcription factor binding sites (TFBSs) based on their z-scores. Z-score was computed based on the frequency of TFBS occurrence detected by the energy method. The z-scores were sorted in ascending order. The TFBSs; CTCF, YY1, and ZNF274, which were previously reported as enrichments in TAD regions are plotted in warm colours. The other TFBSs are plotted in grey colour. (e) The top 10 motifs with the highest z-score on chromosome 10 are listed.

### Predicting TAD boundaries and four sub-categorized TAD boundaries based on digital Hi-C scores

We have shown that our horizontal search approach could predict genomic context based on our prediction results in two following cases: enhancer-promoter interactions and CTCF-enriched regions. Those datasets contained relatively large volumes of positive data which, in general, helped us to conduct better training and accurate outcomes. Next, our challenge was whether the horizontal search approach was still functional to predict genomic contexts even from smaller volumes of positive data.

To address this, we demonstrated to predict TAD boundary regions by applying the horizontal search approach. To conduct this, we again used the datasets of Rao et al., (2014)^20^. In particular, we collected the reference information of TAD boundary regions in IMR90 dataset from the 3D Genome Browser at Northwestern University^48^. We focus on chromosome 10 as a model case. This time, we modified a horizontal search approach in the following two points; using a custom weighted matrix to compute a custom TAD score which was a similar concept of the directionality index^9^, and using a custom TAD score and normalized GC % as input data for RF algorithm (figure 7a). We showed representative results of around 2.98Mbp on chromosome 10. We predicted three regions as TAD boundary regions (i.e., black-colour region) whereas the reference data was called a single TAD boundary region (red-colour region) in the target genomic region. We found only one of our predictions detected reference data; we found that our model returned many ‘false positives’ regions. We closely investigated those regions manually with using the 3D Genome Browser to view raw data. We found that the reference boundary had a strong/clear status whereas one of our predictions showed a weak TAD boundary region. To support this idea, small TAD boundaries were defined (e.g., orange-/blue-color bars) and our prediction was located between the two boundaries (figure 7b). We repeated this process 100 times to compute performance stats. As a result, we found that the model performed 0.766 in accuracy, 0.74 in precision, 0.55 in recall, 0.89 in specificity, and 0.63 in F1 scores (figure 7c). We noticed that the F1 and recall scores were lower than those of the previous cases of the horizontal search approach because the number of defined TAD boundaries was limited (e.g., relatively small number of positive datasets for training). Having said that, the F1 score was 0.63, indicating the RF model and our horizontal search approach were good in the prediction of TAD boundaries as well. In both training and validation datasets were the same, the performance stats on other datasets show similar trends (supplemental figure 8a). When we trained a model on IMR90 and validated the prediction performance on other datasets, we found that some of F1 score were below 0.6 score. However, this might represent that our model could detected weaker TAD boundaries which were not originally defined, computed as negative score, resulting in lower performance stats (supplemental figure 8b).

**Figure 8:**
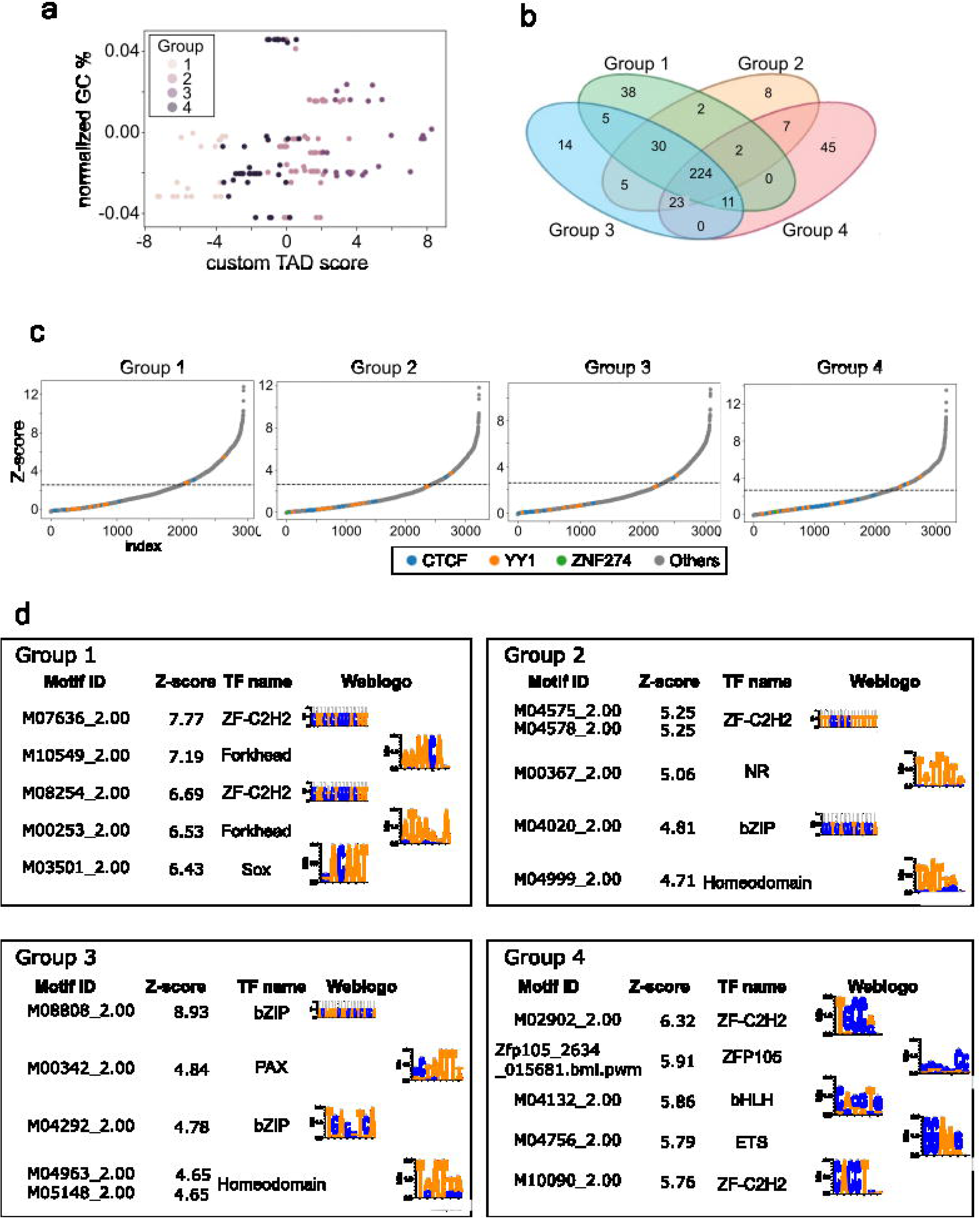
An application of digital Hi-C assay: sub-categorize TAD boundary regions and search the enrichments of motifs in the sub-groups. (a) K-means classification of TAD boundary regions. K = 4 for the K-means algorithm is determined by the Elbow algorithm with custom TAD scores and normalized GC %. (b) Four-way Venn diagram of enriched transcription factor binding sites in TAD boundary regions. (c) Motif search in TAD boundary regions on chromosome 10. Plot transcription factor binding sites (TFBSs) based on their z-scores for individual groups. Z-score was computed based on the frequency of TFBS occurrence detected by the energy method. The z-scores were sorted in ascending order. The TFBSs; CTCF, YY1, and ZNF274, which are previously reported as enrichments in TAD regions are plotted in warm colours. The other TFBSs are plotted in grey colour. (d) The top five motifs with the highest z-score on chromosome 10 are listed per sub- group.

It is known that a couple of transcription factor binding sites (TFBSs) such as CTCF^9^ and YY1^57^ were enriched in the TAD boundary region. Firstly, we addressed whether we could recapture the trend in our data on chromosome 10. To conduct this, we collected 4,719 TFBSs information, (i.e., position weight matrix) or a total of 983 transcription factors from CIS-BP^52^ and UniProbe^53^ (see materials and methods). We predicted binding sites on the boundary regions by applying the energy method^50^. With the same strategy, we randomly selected 500 loci and compute the average occurrence of TFBSs as background stats of the occurrences of TFBSs on the genome. We used those estimated occurrences of each TFBS to compute the z-score of each TFBS for the enrichment of TFBSs in TAD boundary regions. The result was sorted by the z-scores with known enriched TFBSs and showed those regions contained statistical-significant enrichments of known TFBSs; CTCF^9^ (blue-colour dot) and YY1^57^ (orange-colour dot) whereas ZNF274^58^ (green-colour dot) binding sites did not show statistical significancy on chromosome 10 (figure 7d). We found that those known enriched TFBSs did not have the highest score in z-scores. However, we recaptured that the previously known TFBSs, CTCF and YY1, were enriched in TAD boundary regions on chromosome 10. We showed the top 10 TFBSs enrichment with the highest z-scores listed in our prediction results (figure 7e).

Next, we investigated whether these TAD boundary regions could be sub- categorized by using the prediction scores of our horizontal search. We have two reasons. A previous study showed that TAD boundary regions could be sub-categorized into at least two groups, ’strong’ and ’weak’ TAD boundaries, by their binding affinity^47^ under chemical treatments. Hence, we asked whether our digital assay approach also could subcategorise the TAD boundary regions based on our computed scores. Another reason is, while we predicted the TAD boundary regions, we had seen that a trend in reference TAD boundary regions on chromosome 10 could have multiple subgroups by two factors: custom TAD score and normalized GC %. Hence, we addressed this by applying an unsupervised classification method, K-mean, to the scores. We classified the boundary regions into four groups (i.e., K=4) which were determined by the Elbow algorithm (figure 8a). We noted that we could not see the obvious location bias of the members of the subcategorised groups.

Firstly, we investigated the enriched TFBSs in each sub-group. We found each sub- group had the enrichments of known TFBSs (CTCF^9^ and YY1^57^) (figure 8c). When we compared the top 300 enriched TFBSs having the highest z-score in each sub-group (i.e., top 10% of tested TFBSs), the four-way Venn diagram showed that 224 TFBSs were shared across four sub-groups (figure 8b), indicating those boundary regions shared similar or common functions as TAD structures (e.g., structural maintenances). Whereas we found that each group had unique sets of TFBSs, and their sizes were in the range between 8 and 45 TFBSs. For example, Sox (M03501_2.00) in group 1 had 6.43 in z-score and uniquely appeared in this analysis, indicating that those sub-groups might have distinct contributions. We listed that the top five group-specific TFBSs were listed in figure 8d. Additionally, we found that the enriched TFBSs in Groups 1 and 3 remained relatively similar types whereas those in Groups 2 and 4 held similarly. Since that grouping was not reflected based on the score of horizontal searches and GC % (figure 8d), it might exist hidden biological information to characterize the sub-category of TAD boundary regions. Although we needed further experimental and computational verifications, these sub-categories were based on a custom TAD score and GC% which were based on the prediction pattern of our long-distance interactions. Thus, without our predictions, it was hardly sub-categories of the TAD boundary. This result indicates that our digital assay prediction potentially extracts biological features associated with a TAD structure.

In conclusion, we showed that our digital Hi-C assay with vertical and horizontal searches efficiently predicted the long-distance interactions of genomic loci and genomic contexts in three cases: enhancer-promoter interactions, CTCF-enrichments, and TAD-boundary regions. We showed that those trained models could predict these interactions and contexts even in different datasets (different cells/tissues/experimental conditions). Additionally, our prediction scores in our approach could be useful by demonstrating the sub- categorization of the TAD boundaries. Therefore, we have shown that our digital Hi-C assay could efficiently recapture the genomic information from experimental Hi-C assay by applying machine and deep learning methods.

## Discussion

Here, we discuss the following points: generalizing models, the duration time for training, horizontal search, predicting the interaction or active regions, and the advantage of our digital assay.

We ultimately aim to create a more generalized model that describes 3D genome architectures in the status of multiple conditions (e.g., mutants, wild-type). Theoretically, the model allows us to identify unique and descriptive features in a given condition. Identifying those features would be a crucial step to understanding the gene regulatory mechanisms of abnormality or development. Under our current strategy, although the trained models could apply to other tissue/condition datasets, we are still required to create a model per training dataset and chromosome. In our way, we can define key features for a specific condition by detecting the difference between two predicted statuses (e.g., normal, abnormal/mutant). We can also apply the same strategies to the random forest filtering for identifications of PCHi-C and CTCF-enriched regions. In doing so, the current obstacle and limitation is the amount of learning data (e.g., the number of interactions of genomic loci) in training datasets. The mutant/tested data size sometimes can shrink up to 20 folds compared to the control data size. This makes it difficult to fully and accurately learn models, results in a poor prediction rate, overlooks the distinguished interactions and returns high false-positive and true-negative results. To overcome those issues, thus, it is desired to improve our system to satisfy both statuses simultaneously, mutant and wild type.

The long duration time to train each of the models from raw genomic-loci interactions per chromosome is another bottleneck with aiming to develop a more generalized model which describes all human chromosomes. In the performance comparison with other famous deep learning methods, the transformer algorithm does not have to process one word at a time, indicating that the algorithm allows us to parallel than RNNs, thus, reducing training times. However, it still takes a longer time to complete the process in comparison to those LSTM which is often used for traditional sequence-based prediction. We note that the transformer algorithm has an attention process which outperforms LSTM predictions. We maximized the prediction performance of the transformer algorithm to predict long-distance interactions by employing pre-conditions such as the length of the token and input sequences. Those are determined by the threshold of the maximum training duration, 24 hours because the accuracy and loss of the training processes we tested showed saturated results within the cut-off time. With our current set-up (e.g., accessible facility, GPU specs, and programming codes), training a generalized model for all chromosomes will take 24 times longer (i.e., equivalent to more than 24 days for a single dataset). Therefore, it is desired to develop an alternative way (e.g., parallel learning, reducing the volume of training datasets, and developing an efficient algorithm to improve computational speed) to reduce the training models.

Thirdly, we discuss the benefits of pattern analysis of neighbouring target regions (i.e., horizontal search) by applying a random forest (RF) algorithm^54^. Our approach is adapted from a sequence-based approach; hence, the results of our vertical search approach cannot specify whether the regions having predicted/not-predicted are associated with genomic contexts. We complement this shortcoming by developing the horizontal search method for identifying the genomic contexts such as the enhancer-promoter interactions and CTCF-enriched regions. We reasoned that our transformer-basis prediction takes an account of each unit (i.e., 5,000bp-length region) as an independent factor, thus, can handle the bulk information of the neighbouring predictions as new independent information. As a result, we successfully demonstrated that the prediction rates are improved to identify the enhancer-promoter interactions and CTCF-enriched regions at high efficiency. Indeed, our RF models provided a score above 0.8 in F1 scores on validation datasets, indicating that the models are sufficiently good and can recapture the promoter-enhancer interactions in the PCHi-C assay. Those results showed that the horizontal search with the vertical search effectively works to extract the genomic context even by employing the sequence-based approach.

Fourthly, we discuss whether our horizontal search can distinguish the following possibilities: detecting the interactions of genomic loci, the active regulatory regions or both. The regions with long-range interactions correlated with the active/open genomic regions; however, not all active regions associate with long-distance interactions. In our analysis, indeed, our results contain false-positives interactions (i.e., the regions having our predicted interactions but not having captured interactions in the PCHi-C assay). This result may indicate that our horizontal search in the PCHi-C data predicts the active/open regions rather than the enhancer-promoter interactions. Those are speculated under the current models, and it is impossible to distinguish these possibilities. To answer this question, we may conduct experimental verifications with PCHi-C and ATAC-seq.

Lastly, we emphasize the advantages of our digital assay. Due to human and experimental errors, the results of high-throughput assays are often hardly standardized at a high level, resulting in uneven quality across experimental results (e.g., batch effect, rich interactions captured in a dataset but not others). Accuracy and loss scores in our training process could be utilized as quality control; thus, our prediction overcame the quality of the experimental results and maintained high-level prediction rates in true-positive interactions. Additionally, due to financial reasons, researchers tend to have less-duplicated and less-redundant experimental data to collect per condition, which might lose the chance to identify genuine interactions at the end. We demonstrated that our assay could detect more interactions efficiently from a single Hi-C data (figure 3). This result also indicated that our approach could apply a single trained model to other datasets for predicting interactions and have broader applications. We confirmed that these performance stats remained at high levels on other chromosomes (i.e., on chromosomes 5, 10 and 16, in figure 3, supplemental figure 2). Hence, our assay has a general versatility and can cover wider conditions to predict long-distance interactions.

## Supplemental figures

**Supplemental figure 1: Optimization of two factors; the length of the token (six or eight nucleotides) and representative sequences (300 or 600 base pairs)**

Create 10 test-training datasets, each of which contains 1,000,000 interactions selected randomly from the original training dataset (see material and methods) and plot the average scores of accuracy and loss on the test-training datasets in the training process. A cut-off threshold to terminate the training process is 0.99 in accuracy.

(a-f) The transition of (a,d) accuracy, (b,e) loss, and (c, f) time consumption for a single epoch on chromosome 10 up to 50 epochs in the training process. Two factors were tested: the length of the token (six or eight nucleotides) and representative sequences (100, 200, 300 or 600 base pairs).

(g-l) The transition of (g,j) accuracy, (h,k) loss, and (i, l) time consumption for a single epoch on chromosome 20 up to 50 epochs in the training process. Two factors were tested: the length of the token (six or eight nucleotides) and representative sequences (100, 200, 300 or 600 base pairs).

(m-o) The transition of (m) accuracy, (n) loss, and (o) time consumption for a single epoch on chromosome 10 up to 50 epochs in the training process when the length of representative sequences is fixed at 300bp. The variable factor is the length of the token in four (blue), six (orange), and eight (green) nucleotides. The target chromosome is 10.

(p-r) The transition of (p) accuracy, (q) loss, and (r) time consumption for a single epoch on chromosome 10 up to 50 epochs in the training process when the length of representative sequences is fixed at 300bp. The variable factor is the length of the token in four (blue), six (orange), and eight (green) nucleotides. The target chromosome is 20.

**Supplemental figure 2: Visual comparison between predicting and reference interactions of genomic loci and prediction performance statistics Describe the other cases to predict long-distance interactions of genomic loci. On different chromosomes**

(a) A representative stats performance results of predicted interactions with four different reference datasets only (HIC001 - HIC004 from Rao et al., (2014)^20^ in the range of 2,285,000bp to 4,285,000bp on chromosome 10. The location of the input sequence is 3,285,000bp on the chromosome. The “total” (or orange-colour bar) in the last column still indicates the sum of 10 reference scores (HIC001 - HIC010). Prediction performance stats between predicted (positive/negative) and reference (true/false) interactions for each dataset. The performance stats are measured following five factors; accuracy, precision, recall, sensitivity, and F-1 score.

(b) Venn diagram of the predicted and reference interactions at the 200 loci on chromosome 10. The 200 loci are randomly selected on the target chromosome. The model is trained on HIC001. The trained model is used to verify the prediction performance on 10 reference scores (HIC001 - HIC010).

(c) Overall prediction performance stats between predicted (positive/negative) and reference (true/false) interactions at the 200 loci on chromosome 10.

(d) A representative stats performance results of predicted interactions with 4 different reference datasets only (HIC001 - HIC004 from Rao et al., 2014^20^) in the range of 22,505,000 bp to 24,505,000 bp on chromosome 16. The location of the input sequence is 23,505,000 bp on the chromosome. The “total” (or orange-colour bar) in the last column still indicates the sum of 10 reference scores above (HIC001 - HIC010). Prediction performance stats between predicted (positive/negative) and reference (true/false) interactions for each dataset. The performance stats are measured following five factors; accuracy, precision, recall, sensitivity, and F-1 score.

(e) Venn diagram of the predicted and reference interactions at the 200 loci on chromosome

(a) 16. The 200 loci are randomly selected on the target chromosome. The model is trained on HIC001. The trained model is used to verify the prediction performance on 10 reference scores (HIC001 - HIC010).

(b) (f) Overall prediction performance stats between predicted (positive/negative) and reference (true/false) interactions at the 200 loci on chromosome 16.

**Supplemental figure 3: Efficiently predicting the interactions of genomic loci in datasets by using trained models in different datasets.**

(a) Venn diagram of the predicted and reference interactions at the 200 loci on chromosome

(a) 10. The 200 loci are randomly selected on the target chromosome. The model is trained on HIC001. The trained model is used to verify the prediction performance on four reference data (HIC065 - HIC068).

(b) Overall prediction performance stats between predicted (positive/negative) and reference (true/false) interactions at the 200 loci.

(c) A representative stats performance results of predicted interactions with 4 different reference datasets only (HIC065 - HIC068 from Rao et al., 2014) in the range of 32,295,000bp to 34,295,000bp on chromosome 10. The input sequence locates in 33,295,000bp on chromosome 10. The “total” (or orange-colour bar) in the last column still indicates the sum of the four reference scores above (HIC065 - HIC068). Prediction performance stats between predicted (positive/negative) and reference (true/false) interactions for each dataset. The performance stats are measured following five factors; accuracy, precision, recall, sensitivity, and F-1 score.

**Supplemental figure 4: Visual comparison between predicting and reference interactions of smaller-number (N=4) genomic loci and prediction performance statistics**

(a, b) Venn diagram of the predicted and smaller-number (N=4) reference interactions at the 200 loci on chromosome 10. The 200 loci are randomly selected on the target chromosome. The model is trained on HIC001. The trained model is used to verify the prediction performance on two sets of smaller-number reference scores ((a) HIC002 - HIC005, and (b) HIC006 - HIC009).

(c, d) Overall prediction performance stats between predicted (positive/negative) and smaller-number reference (true/false) interactions at the 200 loci on chromosome 10. ((c) HIC002 - HIC005, and (d) HIC006 - HIC009)

(a) (e) A representative stats performance results of predicted interactions with two sets of four different reference datasets only (top: HIC002 - HIC005, bottom: HIC006 - HIC009 from Rao et al., 2014) in the range of 10,000 bp to 2,010,000 bp on chromosome 10. The location of the input sequence is 1,010,000 bp on the chromosome. The orange-colour bar represents the sum of the corresponding four reference scores (i.e., HIC002 - HIC005). Prediction performance stats between predicted (positive/negative) and reference (true/false) interactions for each dataset. The performance stats are measured following five factors; accuracy, precision, recall, sensitivity, and F-1 score. The result in the top row uses HIC002 - HIC005 as reference data while the result in the bottom row uses HIC006 - HIC009.

**Supplemental figure 5: Efficiently predicting the interactions of genomic loci in datasets by using trained models in different datasets (on HIC50, HIC59, and HIC80 on chromosome 10).**

(a-d) Venn diagram of the predicted and reference interactions at the 200 loci on chromosome 10. The 200 loci are randomly selected on the target chromosome. The model is trained on HIC001. The trained model is used to verify the prediction performance on following reference data (a, HIC050 - HIC058, except HIC051; b, HIC059 - HIC065; c, HIC065 - HIC068; d, HIC080 - HIC083 from Rao et al., 2014^20^).

(e-h) Overall prediction performance stats between predicted (positive/negative) and reference (true/false) interactions at the 200 loci on the chromosome. (e, HIC050 - HIC058, except HIC051; f, HIC059 - HIC065; g, HIC065 - HIC068; h, HIC080 - HIC083 from Rao et al., 2014^20^). The performance stats are measured following five factors; accuracy, precision, recall, sensitivity, and F-1 score.

(i-l) A representative stats performance results of predicted interactions with seven different reference datasets (i) HIC050 - HIC058 from Rao et al., 2014 in the range of 18,865,000bp to 20,865,000bp. (j) HIC059 - HIC065 in the range of 33,145,000bp to 35,145,000bp. (k) HIC065 - HIC068 in the range of 32,295,000bp to 34,295,000bp. (l) HIC080 - HIC083 in the range of 32,295,000bp to 34,295,000bp. The input sequence locates in the mid-position in each target region. The performance stats are measured following five factors; accuracy, precision, recall, sensitivity, and F-1 score.

**Supplemental figure 6: Prediction performance statistics on other tissue types for promoter-capture Hi-C.**

(a-aa) Venn diagrams for the overlap of predicted and various kinds of reference data and Overall prediction performance stats between AO-model prediction (positive/negative) and various kinds of reference (true/false) interactions at the 200 loci on chromosome 10. The 200 loci are randomly selected on the target chromosome. The model was created on the AO data as a model case. The developed model was used to verify the prediction performance on the following reference data; (a) AD, (b) CM, (c) DLPFC, (d) EG, (e) FT, (f) GA, (g) GM, (h) H1, (i) HC, (j) IMR90, (k) LG, (l) LI, (m) LV, (n) ME, (o) MSC, (p) NPC, (q) OV, (r) PA, (s) PO, (t) RA, (u) RV, (v) SB, (w) SG, (x) SX, (y) TB, (z) TH, and (aa) ALL. ‘ALL’ contained all interactions across 27 tissues in PCHi-C datasets.

**Supplemental figure 7: Predictions of CTCF-enriched regions**

(a, b, e, f) Venn diagram for the overlap of predicted and reference data at the 200 loci on (a, b) chromosome 5 and on (e, f) chromosome 16. The 200 loci are randomly selected on each target chromosome. The model was created on the training data and validated their performance on validation datasets with the following combinations: (a, e) the training data are LNCaP cell data, and the validation data are IMR90 cell data while (b, f) the training data are IMR90 cell data and the validation data are LNCaP cell data.

(c, d, g, h) Overall prediction performance stats at the 200 loci on (c, d) chromosome 5 and on (g, h) chromosome 16. (c, g) has the training data are LNCaP cell data and the validation data are IMR90 cell data while (d, h) has the training data are IMR90 cell data and the validation data are LNCaP cell data

**Supplemental figure 8: Predictions of TAD-boundary regions:**

(a) Comparison of prediction performance stats across different tissues on chromosome 10. The target chromosome is chromosome 10. Each dataset splits into training (90%) and validation (10%) datasets with shuffling. Each time, performance stats are computed for the following five factors; accuracy, precision, recall, sensitivity, and F-1 score. Repeat this process 100 times.
(b) Cross-validation of prediction performance stats across different tissues on chromosome 10. Each model was trained on the IMR90 dataset and validated its prediction performance on other six datasets (GM12878, HMEC, HUVEC, K562, KBM7, NHEK). Each time, performance stats are computed for the following five factors; accuracy, precision, recall, sensitivity, and F-1 score. Repeat this process 100 times.

## Acknowledgements

The authors are grateful to the members of the GS lab for the fruitful discussion. AM is thankful to Dr Natasha Naumova for the valuable discussion on initiating this project.

## Author contributions

AM conceived the project, and designed, conducted and analysed computational works. AM wrote the manuscript and finalized it with GS’s help.

## Reference

1. Blackwood, E. M. & Kadonaga, J. T. Going the distance: a current view of enhancer action. Science 281, 60–63 (1998).

2. Williamson, I., Hill, R. E. & Bickmore, W. A. Enhancers: from developmental genetics to the genetics of common human disease. Dev. Cell 21, 17–19 (2011).

3. Pennacchio, L. A., Bickmore, W., Dean, A., Nobrega, M. A. & Bejerano, G. Enhancers: five essential questions. Nature reviews. Genetics vol. 14 288–295 (2013).

4. Long, H. K. et al. Loss of Extreme Long-Range Enhancers in Human Neural Crest Drives a Craniofacial Disorder. Cell Stem Cell 27, 765–783.e14 (2020).

5. Sakabe, N. J., Savic, D. & Nobrega, M. A. Transcriptional enhancers in development and disease. Genome Biol. 13, 238 (2012).

6. Ryan, G. E. & Farley, E. K. Functional genomic approaches to elucidate the role of enhancers during development. Wiley Interdiscip. Rev. Syst. Biol. Med. 12, e1467 (2020).

7. Hindorff, L. A. et al. Potential etiologic and functional implications of genome- wide association loci for human diseases and traits. Proc. Natl. Acad. Sci. U. S. A. 106, 9362–9367 (2009).

8. Maurano, M. T. et al. Systematic localization of common disease-associated variation in regulatory DNA. Science 337, 1190–1195 (2012).

9. Dixon, J. R. et al. Topological domains in mammalian genomes identified by analysis of chromatin interactions. Nature 485, 376–380 (2012).

10. Boyle, A. P. et al. High-resolution mapping and characterization of open chromatin across the genome. Cell 132, 311–322 (2008).

11. Giresi, P. G., Kim, J., McDaniell, R. M., Iyer, V. R. & Lieb, J. D. FAIRE (Formaldehyde-Assisted Isolation of Regulatory Elements) isolates active regulatory elements from human chromatin. Genome Res. 17, 877–885 (2007).

12. Buenrostro, J. D., Giresi, P. G., Zaba, L. C., Chang, H. Y. & Greenleaf, W. J. Transposition of native chromatin for fast and sensitive epigenomic profiling of open chromatin, DNA-binding proteins and nucleosome position. Nat. Methods 10, 1213–1218 (2013).

13. Johnson, D. S., Mortazavi, A., Myers, R. M. & Wold, B. Genome-wide mapping of in vivo protein-DNA interactions. Science 316, 1497–1502 (2007).

14. Lieberman-Aiden, E. et al. Comprehensive mapping of long-range interactions reveals folding principles of the human genome. Science 326, 289–293 (2009).

15. Fullwood, M. J. et al. An oestrogen-receptor-alpha-bound human chromatin interactome. Nature 462, 58–64 (2009).

16. Tang, Z. et al. CTCF-Mediated Human 3D Genome Architecture Reveals Chromatin Topology for Transcription. Cell 163, 1611–1627 (2015).

17. Dekker, J., Rippe, K., Dekker, M. & Kleckner, N. Capturing chromosome conformation. Science 295, 1306–1311 (2002).

18. Schoenfelder, S., Javierre, B.-M., Furlan-Magaril, M., Wingett, S. W. & Fraser, P. Promoter Capture Hi-C: High-resolution, Genome-wide Profiling of Promoter Interactions. J. Vis. Exp. (2018) doi:10.3791/57320.

19. Jung, I. et al. A compendium of promoter-centered long-range chromatin interactions in the human genome. Nat. Genet. 51, 1442–1449 (2019).

20. Rao, S. S. P. et al. A 3D map of the human genome at kilobase resolution reveals principles of chromatin looping. Cell 159, 1665–1680 (2014).

21. Hsieh, T.-H. S. et al. Mapping Nucleosome Resolution Chromosome Folding in Yeast by Micro-C. Cell 162, 108–119 (2015).

22. Freire-Pritchett, P. et al. Detecting chromosomal interactions in Capture Hi-C data with CHiCAGO and companion tools. Nat. Protoc. 16, 4144–4176 (2021).

23. Zeng, W., Wu, M. & Jiang, R. Prediction of enhancer-promoter interactions via natural language processing. BMC Genomics 19, 84 (2018).

24. Ron, G., Globerson, Y., Moran, D. & Kaplan, T. Promoter-enhancer interactions identified from Hi-C data using probabilistic models and hierarchical topological domains. Nat. Commun. 8, 2237 (2017).

25. Singh, S., Yang, Y., Póczos, B. & Ma, J. Predicting enhancer-promoter interaction from genomic sequence with deep neural networks. Quant. Biol. (Beijing, China) 7, 122–137 (2019).

26. Avsec, Ž., et al. Effective gene expression prediction from sequence by integrating long-range interactions. Nat. Methods 18, 1196–1203 (2021).

27. Zhou, J. et al. Deep learning sequence-based ab initio prediction of variant effects on expression and disease risk. Nat. Genet. 50, 1171–1179 (2018).

28. Kelley, D. R. Cross-species regulatory sequence activity prediction. PLoS Comput. Biol. 16, e1008050 (2020).

29. Roy, S. et al. A predictive modeling approach for cell line-specific long-range regulatory interactions. Nucleic Acids Res. 43, 8694–8712 (2015).

30. Hait, T. A., Amar, D., Shamir, R. & Elkon, R. FOCS: A novel method for analyzing enhancer and gene activity patterns infers an extensive enhancer- promoter map. Genome Biol. 19, 1–14 (2018).

31. Gao, T. & Qian, J. Eagle: An algorithm that utilizes a small number of genomic features to predict tissue/ cell type-specific enhancer-gene interactions. PLoS Comput. Biol. 15, 1–22 (2019).

32. Liu, Y., Barr, K. & Reinitz, J. Fully interpretable deep learning model of transcriptional control. Bioinformatics 36, i499–i507 (2020).

33. Whalen, S., Truty, R. M. & Pollard, K. S. Enhancer-promoter interactions are encoded by complex genomic signatures on looping chromatin. Nat. Genet. 48, 488–496 (2016).

34. Mills, C. et al. PEREGRINE: A genome-wide prediction of enhancer to gene relationships supported by experimental evidence. PLoS One 15, 1–22 (2020).

35. Kelley, D. R. et al. Sequential regulatory activity prediction across chromosomes with convolutional neural networks. Genome Res. 28, 739–750 (2018).

36. Wang, H., Huang, B. & Wang, J. Predict long-range enhancer regulation based on protein-protein interactions between transcription factors. Nucleic Acids Res. 49, 10347–10368 (2021).

37. Zhuang, Z., Shen, X. & Pan, W. A simple convolutional neural network for prediction of enhancer-promoter interactions with DNA sequence data. Bioinformatics 35, 2899–2906 (2019).

38. Mao, W., Kostka, D. & Chikina, M. Modeling Enhancer-Promoter Interactions with Attention-Based Neural Networks. bioRxiv 219667 (2017).

39. Umarov, R. et al. ReFeaFi: Genome-wide prediction of regulatory elements driving transcription initiation. PLoS Comput. Biol. 17, 1–18 (2021).

40. Zhang, M., Hu, Y. & Zhu, M. EPIshilbert: Prediction of enhancer-promoter interactions via hilbert curve encoding and transfer learning. Genes (Basel). 12, (2021).

41. Schwessinger, R. et al. DeepC: predicting 3D genome folding using megabase-scale transfer learning. Nat. Methods 17, 1118–1124 (2020).

42. Hong, Z., Zeng, X., Wei, L. & Liu, X. Identifying enhancer-promoter interactions with neural network based on pre-trained DNA vectors and attention mechanism. Bioinformatics 36, 1037–1043 (2020).

43. Belokopytova, P. S., Nuriddinov, M. A., Mozheiko, E. A., Fishman, D. & Fishman, V. Quantitative prediction of enhancer-promoter interactions. Genome Res. 30, 72–84 (2020).

44. Vaswani, A. et al. Attention is all you need. Adv. Neural Inf. Process. Syst. 2017-Decem, 5999–6009 (2017).

45. Hochreiter, S. & Schmidhuber, J. Long Short-Term Memory. Neural Comput. 9, 1735–1780 (1997).

46. Jung, I. et al. A compendium of promoter-centered long-range chromatin interactions in the human genome. Nat. Genet. 51, 1442–1449 (2019).

47. Khoury, A. et al. Constitutively bound CTCF sites maintain 3D chromatin architecture and long-range epigenetically regulated domains. Nat. Commun. 11, (2020).

48. Wang, Y. et al. The 3D Genome Browser: a web-based browser for visualizing 3D genome organization and long-range chromatin interactions. Genome Biol. 19, 151 (2018).

49. Knight, P. A. & Ruiz, D. A fast algorithm for matrix balancing. IMA J. Numer. Anal. 33, 1029–1047 (2013).

50. Zhao, Y., Granas, D. & Stormo, G. D. Inferring binding energies from selected binding sites. PLoS Comput. Biol. (2009) doi:10.1371/journal.pcbi.1000590.

51. Fuxman Bass, J. I., et al. Human gene-centered transcription factor networks for enhancers and disease variants. Cell 161, 661–673 (2015).

52. Weirauch, M. T. et al. Determination and Inference of Eukaryotic Transcription Factor Sequence Specificity. Cell 158, 1431–1443 (2014).

53. Hume, M. A., Barrera, L. A., Gisselbrecht, S. S. & Bulyk, M. L. UniPROBE, update 2015: New tools and content for the online database of protein-binding microarray data on protein-DNA interactions. Nucleic Acids Res. 43, D117– D122 (2015).

54. Tin Kam Ho. Random Decision Forests Tin Kam Ho Perceptron training. Proc. 3rd Int. Conf. Doc. Anal. Recognit. 278–282 (1995).

55. Singh, S. et al. Predicting Enhancer-Promoter Interaction from Genomic Sequence with Deep Learning. bioRxiv 7, 1–5 (2018).

56. Van Bortle, K. et al. Insulator function and topological domain border strength scale with architectural protein occupancy. Genome Biol. 15, R82 (2014).

57. Weintraub, A. S. et al. YY1 Is a Structural Regulator of Enhancer-Promoter Loops. Cell 171, 1573–1588.e28 (2017).

58. Hong, S. & Kim, D. Computational characterization of chromatin domain boundary-associated genomic elements. Nucleic Acids Res. 45, 10403–10414 (2017).

