## Supplementary figures and images for "Development of digital Hi-C assay"

### supplemental figure 1

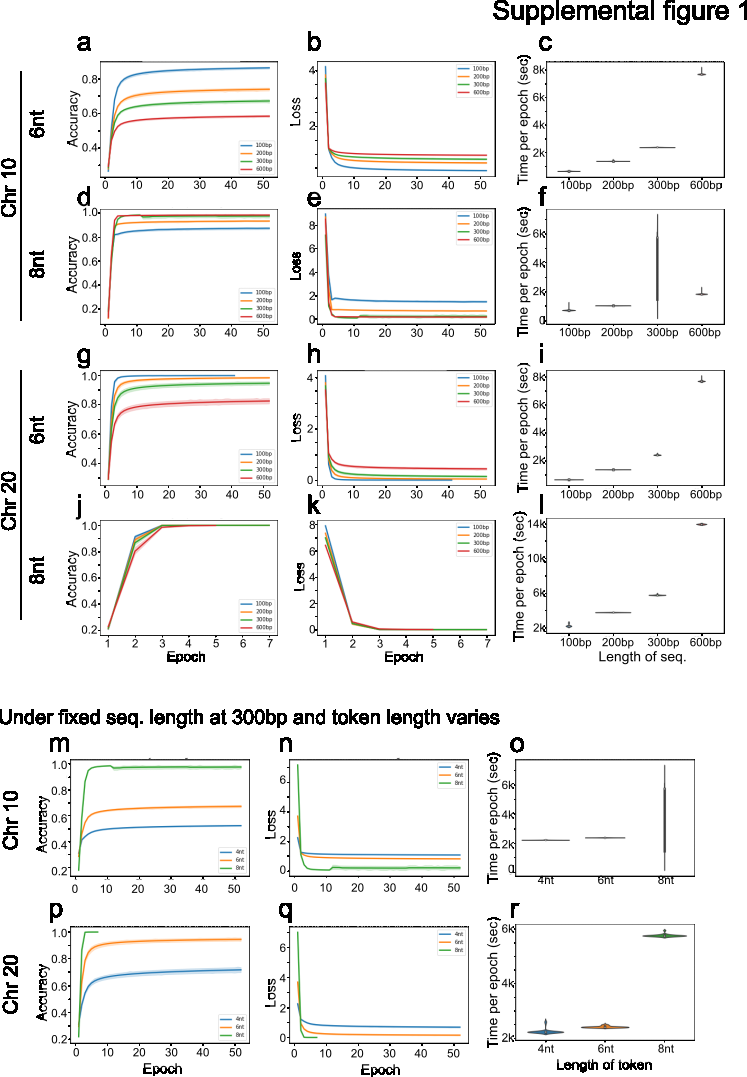

### supplemental figure 2

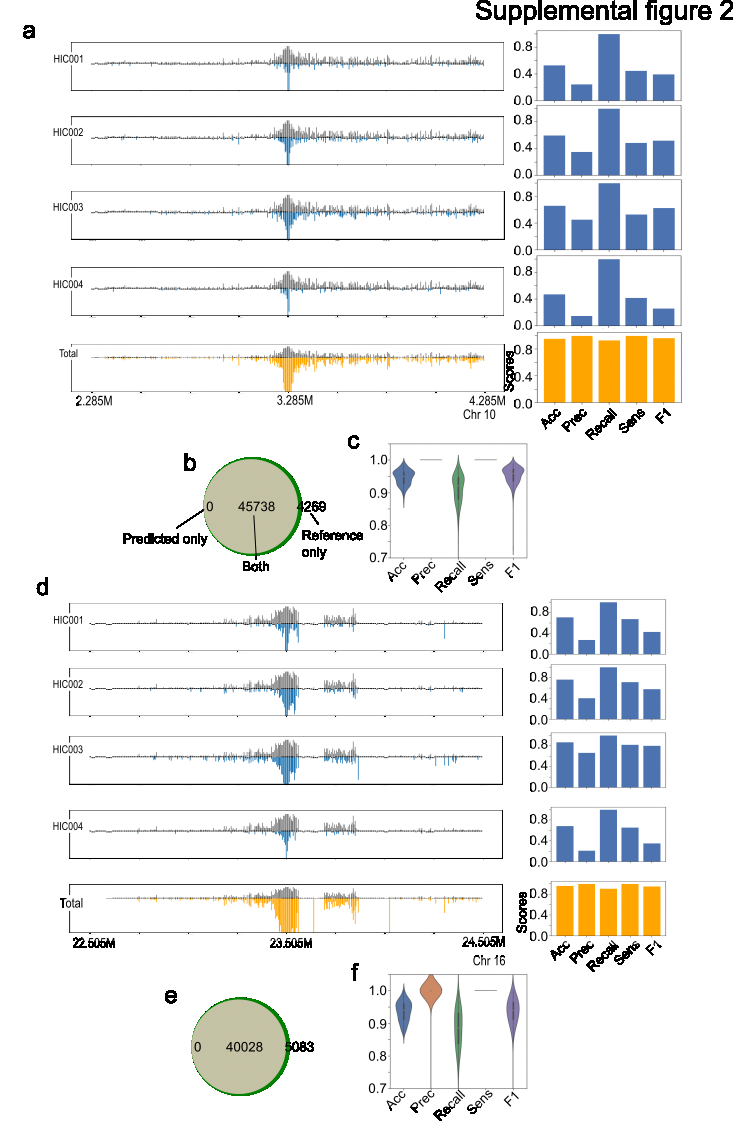

### supplemental figure 3

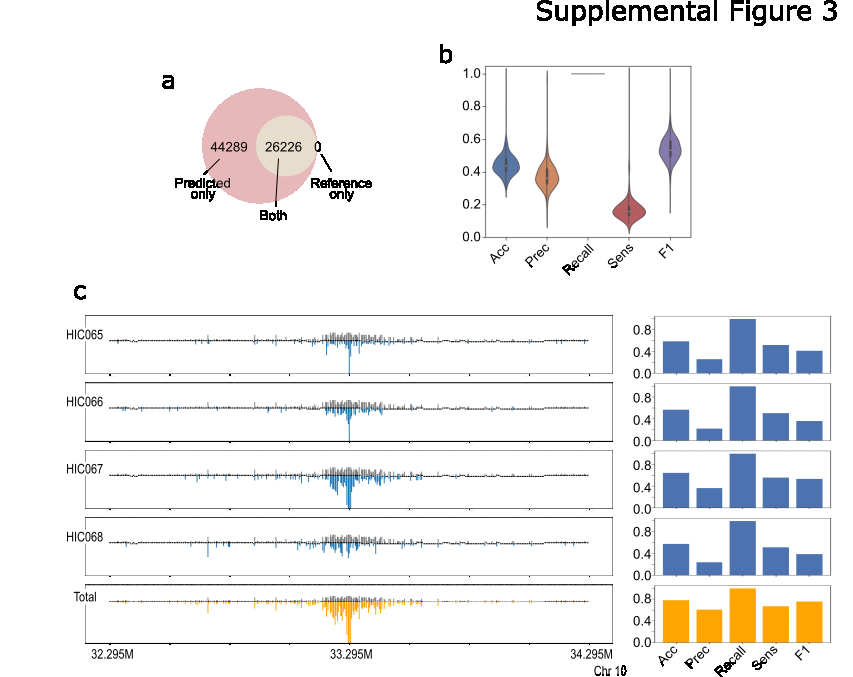

### supplemental figure 4

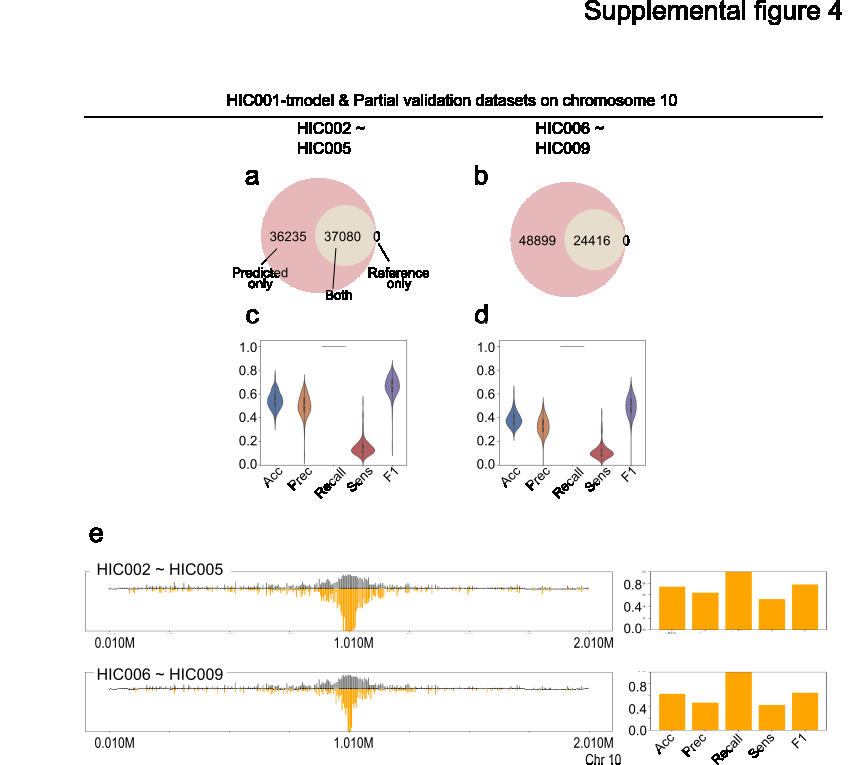

### supplemental figure 5

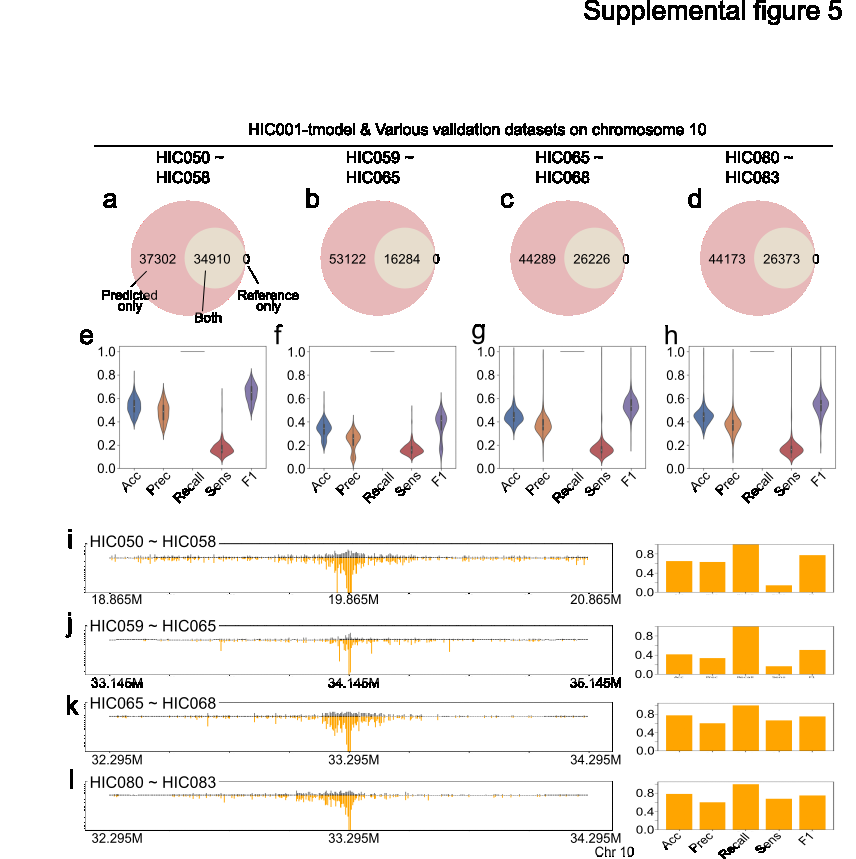

### supplemental figure 6

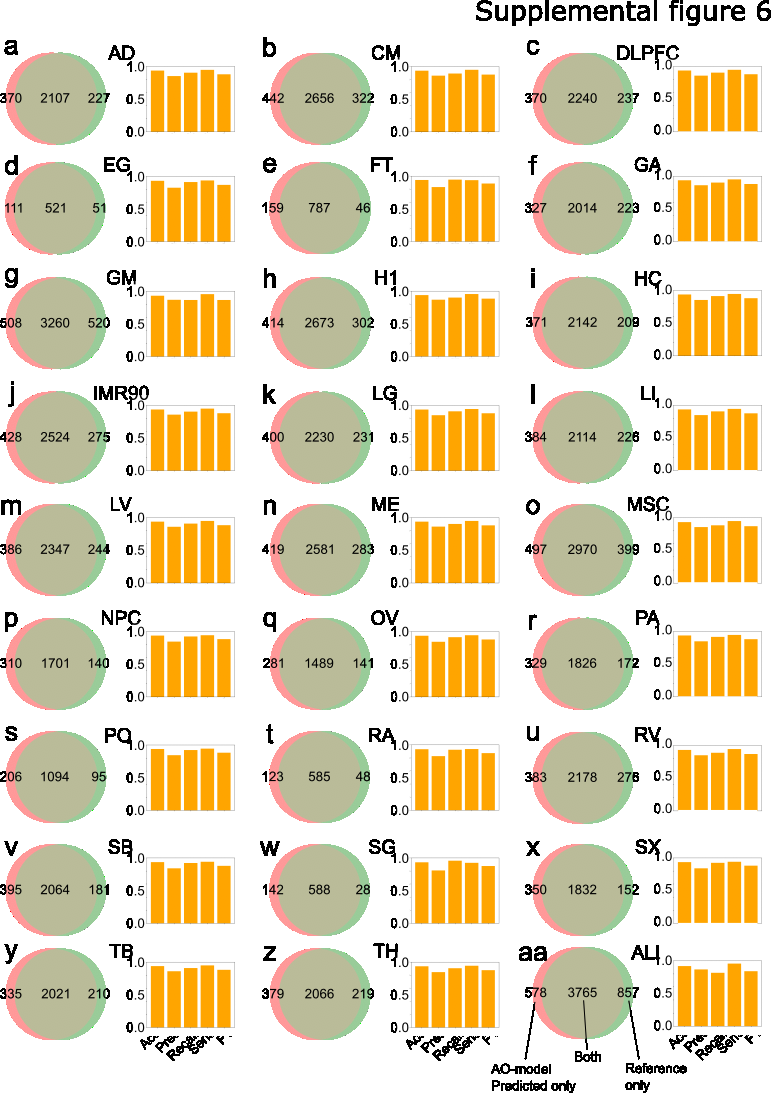

### supplemental figure 7

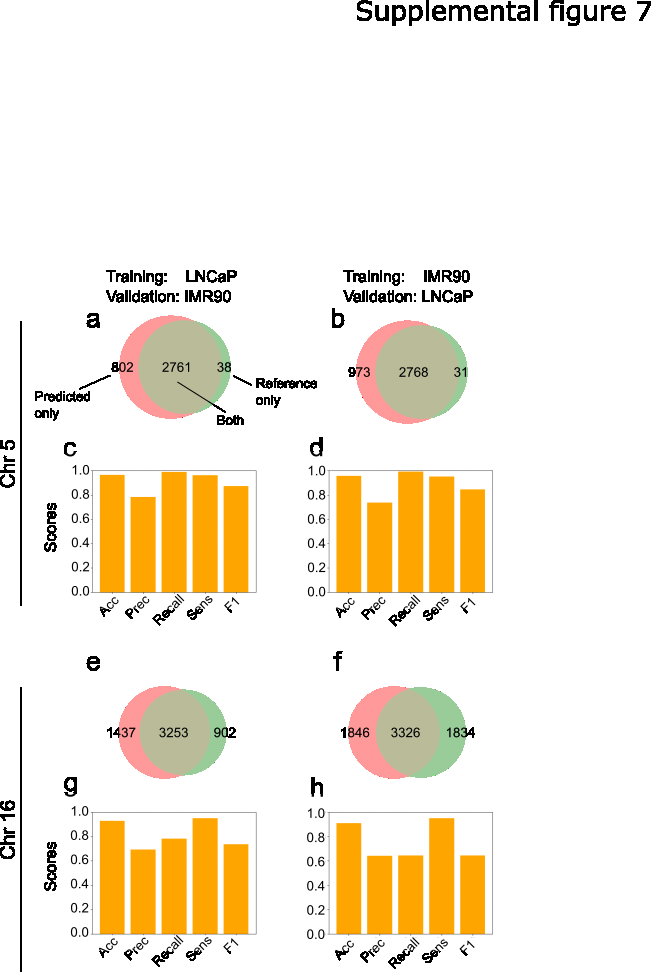

### supplemental figure 8

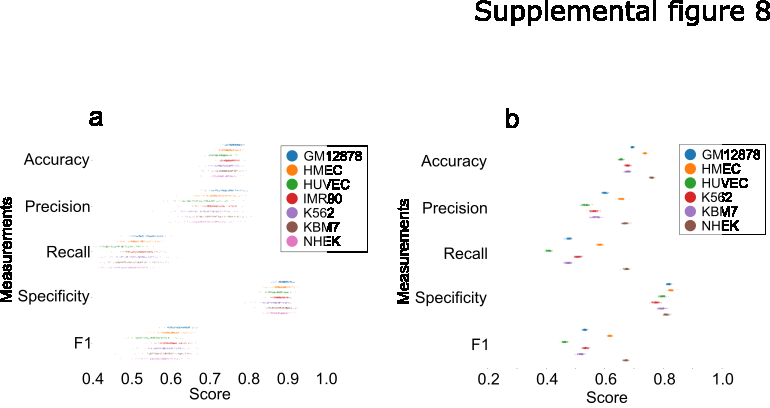
